# Immunogenicity and protective efficacy of a highly thermotolerant, trimeric SARS-CoV-2 receptor binding domain derivative

**DOI:** 10.1101/2021.01.13.426626

**Authors:** Sameer Kumar Malladi, Unnatiben Rajeshbhai Patel, Raju S Rajmani, Randhir Singh, Suman Pandey, Sahil Kumar, Sara Khaleeq, Petrus Jansen van Vuren, Shane Riddell, Sarah Goldie, Savitha Gayathri, Debajyoti Chakraborty, Parismita Kalita, Ishika Pramanick, Nupur Agarwal, Poorvi Reddy, Nidhi Girish, Aditya Upadhyaya, Mohammad Suhail Khan, Kawkab Kanjo, Madhuraj Bhat, Shailendra Mani, Sankar Bhattacharyya, Samreen Siddiqui, Akansha Tyagi, Sujeet Jha, Rajesh Pandey, Shashank Tripathi, Somnath Dutta, Alexander J. McAuley, Nagendrakumar Balasubramanian Singanallur, Seshadri S. Vasan, Rajesh P. Ringe, Raghavan Varadarajan

## Abstract

The Receptor Binding Domain (RBD) of SARS-CoV-2 is the primary target of neutralizing antibodies. We designed a trimeric, highly thermotolerant glycan engineered RBD by fusion to a heterologous, poorly immunogenic disulfide linked trimerization domain derived from cartilage matrix protein. The protein expressed at a yield of ∼80-100 mg/liter in transiently transfected Expi293 cells, as well as CHO and HEK293 stable cell lines and formed homogeneous disulfide-linked trimers. When lyophilized, these possessed remarkable functional stability to transient thermal stress of upto 100 °C and were stable to long term storage of over 4 weeks at 37 °C unlike an alternative RBD-trimer with a different trimerization domain. Two intramuscular immunizations with a human-compatible SWE adjuvanted formulation, elicited antibodies with pseudoviral neutralizing titers in guinea pigs and mice that were 25-250 fold higher than corresponding values in human convalescent sera. Against the beta (B.1.351) variant of concern (VOC), pseudoviral neutralization titers for RBD trimer were ∼ three-fold lower than against wildtype B.1 virus. RBD was also displayed on a designed ferritin-like Msdps2 nanoparticle. This showed decreased yield and immunogenicity relative to trimeric RBD. Replicative virus neutralization assays using mouse sera demonstrated that antibodies induced by the trimers neutralized all four VOC to date, namely B.1.1.7, B.1.351, P.1 and B.1.617.2 without significant differences. Trimeric RBD immunized hamsters were protected from viral challenge. The excellent immunogenicity, thermotolerance, and high yield of these immunogens suggest that they are a promising modality to combat COVID-19, including all SARS-CoV-2 VOC to date.

---

The Coronavirus infectious disease 2019 (COVID-19) pandemic caused by SARS-CoV-2^1,2^ has led to ∼177.1 million infections and ∼3.8 million deaths worldwide as on 21^st^ June, 2021^3^. India experienced a debilitating second wave, with one of the highest daily infection rates in the world. The viral spike glycoprotein is the most abundant protein exposed on the viral surface and the primary target of host elicited humoral immune responses^4–15^. Thus, there are a large number of COVID-19 vaccine candidates in various stages of development, with ∼ 11 candidates already granted emergency use authorisation. However, all of these are required to be stored either refrigerated or frozen. There is thus an unmet need for efficacious vaccines that can be stored for extended periods at room temperature. In addition, there are recent reports of new strains of the virus with enhanced transmissibility and immune evasion^16,17^. This emphasizes the urgent need to develop vaccine formulations that elicit high titers of neutralizing antibodies to buffer against viral sequence variation^18–20^. Spike glycoprotein, like various Class I viral surface glycoproteins, assembles as a trimer with each protomer composed of the surface exposed S1 and membrane anchored S2 subunit^21^. The S1 subunit consists of four independently folding domains: N-terminal domain (NTD), Receptor binding domain (RBD), and two short domains (SD1 and SD2) connected by linker regions^4,5,22^. The receptor binding domain (RBD) contains the receptor binding motif (residues 438-505) that facilitates interaction with the ACE2 receptor. Subsequent fusion or endocytosis is mediated by the fusion peptide that constitutes the N-terminal stretch of the S2 subunit^21^. It is now well understood that the majority of neutralizing antibodies in both natural infection and vaccination target the RBD^8,9,11,12,23–28^. Thus, various groups are involved in designing RBD-based immunogens^29–40^. We have previously designed a glycan engineered RBD derivative that was highly thermotolerant and induced moderate titers of neutralizing antibodies^37^. Monomeric versions of immunogens elicit lower binding and neutralizing antibodies than multimeric versions^29,37,40,41^. Potential strategies to improve neutralizing antibody titers include fusions containing repetitive antigenic proteins, Fc fusion based dimerization, nanoparticle design and display strategies, and VLP based display platforms^31,32,38–43^. While effective, several display strategies lead to significant antibody titers against the display scaffold or oligomerization motif, such antibodies might either show undesirable side effects in a small fraction of individuals or direct the response away from the intended target after repeated immunizations. In an alternative strategy, we fused our previously described thermotolerant RBD^37^ to a trimerization motif, namely a disulfide linked coiled-coil trimerization domain derived from human cartilage matrix protein (hCMP), to the N-terminus of mRBD. This trimerization domain is expected to be less immunogenic in small animals due to its homology with the corresponding ortholog, than other widely used trimerization domains of bacterial or synthetic origin such as foldon or IZ^44^. In order to compare trimeric RBD with nanoparticle displayed RBD, we also displayed RBD on the surface of ferritin like nanoparticles, employing SpyCatcher-SpyTag technology^45,46^. hCMP-mRBD expressed as homogenous trimers, possessed comparable thermal stability profiles to the corresponding monomer^37^ and remained functional after over 4 weeks upon lyophilization and storage at 37 °C. The trimeric RBD is highly immunogenic in mice and guinea pigs when formulated with SWE adjuvant. SWE is equivalent to the widely used, clinically approved, MF59 adjuvant^47^. Oligomerization increased neutralizing antibody titers by ∼25-250 fold when compared with the titers in human convalescent sera, providing a proof of principle for the design strategy. Further the hCMP-mRBD protected hamsters from viral challenge, and immunized sera from mice and guinea pigs neutralized the rapidly spreading B.1.351 viral variant with only a three-fold decrease in neutralization titers. Stable *CHO* and *HEK293* cell lines expressing hCMP-mRBD were constructed and the corresponding protein was as immunogenic, as the protein expressed from transient transfection. Nanoparticle displayed RBD was expressed at lower yield and did not confer any apparent advantage in immunogenicity relative to trimeric RBD. The very high thermotolerance, enhanced immunogenicity, and protection from viral challenge suggest that this trimeric mRBD with inter-subunit, stable disulfides, is an attractive vaccine candidate that can be deployed to combat COVID-19 without requirement of a cold-chain, especially in resource limited settings.

## Results

### Design of trimeric RBDs of SARS-CoV-2

We previously designed a monomeric glycan engineered derivative of the receptor binding domain termed mRBD (residues 332-532 possessing an additional glycosylation site at N532) that induced neutralizing antibodies in guinea pig immunizations^37^. It is known that oligomerization of native antigens can induce higher titers of binding and neutralizing antibodies^31,40,42,48–52^. We therefore fused mRBD to the disulfide linked trimerization domain derived from human cartilage matrix protein (hCMP) (residues 298-340). We have previously used this domain to successfully trimerize derivatives of HIV-1 gp120. These earlier derivatives were used to successfully elicit high titers of broadly reactive anti-gp120 antibodies in guinea pigs, and rabbits. In rhesus macaques when combined with an MVA prime, the formulation conferred protection against heterologous SHIV challenge, without apparent adverse effects^53–55^. We hypothesized that RBD fused to the hCMP trimerization domain (residues 298-340), would elicit higher neutralizing antibody titers relative to the corresponding monomer. In the closed state structure model of Spike-2P protein (PDB 6VXX, residues 332-532), the three RBDs are in the down conformation. We separated the coaxially aligned hCMP trimerization domain C-terminal residue 340, Cα plane from the RBD N-terminal Cα plane by ∼22 Å to eliminate any steric clashes (Figure 1a). The distance between the hCMP C-terminus residue 340 and RBD N-terminus residue 332 was ∼ 39.0 Å in the modeled structure (Figure 1a). A fourteen amino acid linker L14 will comfortably span this distance. We employed the same trimerization domain-linker combination used in our previously described HIV-1 gp120 trimer design^56^. Thus, the trimeric hCMP-mRBD design consisted of the N-terminal hCMP trimeric coiled coil domain (residues 298-340) fused to the I332 residue of mRBD by the above linker, followed by the cleavable His tag sequence described previously^37^ (Figure 1b). The hCMP trimerization domain leads to formation of covalently stabilized trimers crosslinked by interchain disulfides in the hCMP domain. This design is termed hCMP-mRBD and hCMP pRBD where the “m” and “p” signifies expression in mammalian or *Pichia pastoris* cells, respectively.

**Figure 1.**
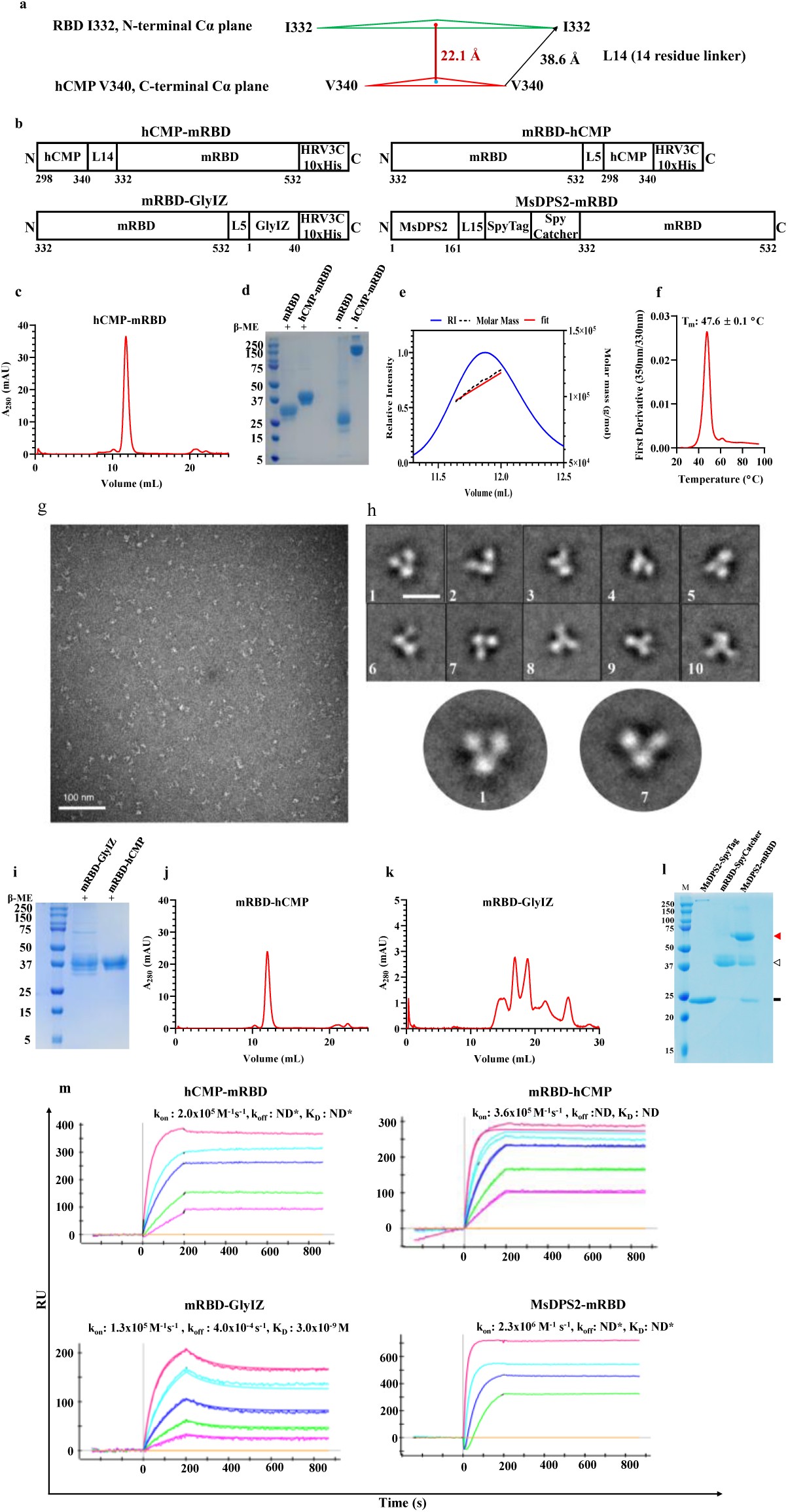
Design and characterization of trimeric mRBD. **a.** The design utilized the RBD (residues 332-532) from the closed state of the Spike-2P (PDB 6VXX) aligned coaxially withthe hCMP trimerization domain, coordinates taken from the homolog CCMP (PDB:1AQ5, Chain 1.1). The N termini of mRBD are labelled as I332 and the hCMP trimerization domain C-termini are labelled as V340. The N, C termini Cα’s form vertices of equilateral triangles. The N-terminal plane of RBD (I332) is separated from the C-terminal plane (V340) of the hCMP trimerization domain by ∼22.1 Å to avoid steric clashes. The I332 terminus and V340terminus are ∼39 Å apart in the modelled structure and are connected by a 14-residue long linker. **b.** hCMP-mRBD consists of N-terminal hCMP trimerization domain fused to I332 of RBD by a linker (L14). mRBD-hCMP consists of the C-terminal hCMP trimerization domainfused to N532 of RBD by a linker (L5). mRBD-GlyIZ consists of a C-terminal GlyIZtrimerization domain fused to N532 of RBD by a linker (L5). MsDPS2-mRBD consists of theMsDPS2 nanoparticle fused to SpyTag covalently linked with mRBD-SpyCatcher. **c.** SEC elution profile of trimeric hCMP-mRBD. **d.** SDS-PAGE of purified mRBD and hCMP-mRBD in reducing and non-reducing conditions demonstrating formation of disulfide-linked trimers. **e.** SEC-MALS of purified hCMP-mRBD (MW: 110 ±10 kDa). The red, black and blue profilesare of the molar mass fit, molar mass and refractive index (RI) respectively. **f.** nanoDSF equilibrium thermal unfolding of hCMP-mRBD. **g**. A representative negative staining image of hCMP-RBD protein. **h**. Representative reference free 2D class averages of hCMP-RBD. 2D class averages indicate that the hCMP-RBD protein is monodisperse and stable. The protein forms a stable trimer. The scale bar shown in the 2D class average is 20 nm. The bottom panel shows the enlarged view of 2D class averages, which specify trimeric hCMP-RBD protein. **i.** SDS-PAGE of purified mRBD-GlyIZ and mRBD-hCMP in reducing conditions. **j, k.** SEC elution profiles of mRBD-hCMP (j) and mRBD-GlyIZ (k). **l.** SDS-PAGE of purified MsDpS2-SpyTag, mRBD-SpyCatcher and the resulting MsDPS2-SpyTag-mRBD-SpyCatcher conjugate abbreviated MsDPS2-mRBD for simplicity. The black solid line, triangle without fill and red triangle correspond to MsDPS2-SpyTag nanoparticle, mRDS-SpyCatcher and MsDPS2-mRBD conjugate respectively. **m.** SPR binding of hCMP-mRBD, mRBD-hCMP, mRBD-GlyIZ and SEC purified complex MsDPS2-mRBD to immobilized ACE2. The curves from highest to lowest correspond to concentrations100 nM, 50 nM, 25 nM, 12.5 nM and 6.25 nM respectively for hCMP-mRBD, mRBD-hCMPand mRBD-GlyIZ. The curves for MsDPS2-mRBD correspond from highest to lowest concentrations of 10 nM, 5 nM, 2.5 nM and 1.25 nM respectively. ND*- No dissociation.

Further, we also designed trimeric RBD constructs (residues 332-532) by fusing hCMP and glycosylated IZ^44^ synthetic trimerization domains at the C-terminus of RBD (Figure 1b). GlyIZ is a glycosylated version of the synthetic trimerization domain IZ. The glycosylation results in immunosilencing of the otherwise highly immunogenic IZ sequence^44^. These constructs were named mRBD-hCMP and mRBD-GlyIZ respectively (Figure 1b). Additionally, we constructed a SpyCatcher^45^ fusion of mRBD by fusion of SpyCatcher^45^ to the C-terminus of the mRBD. This construct is referred to as mRBD-SpyCatcher. These fusion constructs are expressed from transiently transfected mammalian cell culture platforms. A dodecameric self-assembling nanoparticle (MsDPS2) from *Mycobacterium smegmatis* was fused to SpyTag^45,46^ by a 15 residue linker to aid in the complexation of nanoparticle with mRBD-SpyCatcher (Figure 1b). We have successfully employed the MsDPS2 nanoparticle to display trimeric influenza stem immunogens^57^.

### hCMP-mRBD forms homogenous, thermotolerant trimers

hCMP-mRBD was first expressed by transient transfection in Expi293F suspension cells, followed by single step metal affinity chromatography (Ni-NTA) and tag cleavage. The purified protein was observed to be pure and trimeric by reducing and non-reducing SDS-PAGE (Figure 1c, 1d). The protein exists as a homogenous trimer in solution and the molar mass was determined by SEC-MALS to be 110 ±10 kDa, consistent with the presence of nine glycosylation sites in the trimer (Figure 1c, 1e). Trimeric hCMP-mRBD was observed to have comparable thermal stability (T_m_: 47.6 °C) as monomeric mRBD (T_m_: 50.3 °C) (Figure 1f), negative stain EM analysis confirmed the trilobed arrangement of hCMP-mRBD structure (Figure 1g, 1h and Supporting Figure S1) and trimeric RBD bound both its cognate receptor ACE2 and a SARS-CoV-1 neutralizing antibody CR3022 with very high affinity (K_D_ <1nM) and negligible dissociation (Figure 1m, Supporting Figure S2).

The fusion constructs mRBD-hCMP and mRBD-GlyIZ were purified from transiently transfected *Expi293F* cells. mRBD-GlyIZ was observed to be more heterogeneous compared to hCMP-mRBD and mRBD-hCMP (Figure 1c, 1i, 1j, 1k). mRBD-hCMP showed negligible dissociation and bound ACE2 and CR3022 similar to hCMP-mRBD (Figure 1m, Supporting Figure S2). mRBD-GlyIZ bound ACE2 and CR3022 with a K_D_ of 3-5 nM (Figure 1m, Supporting Figure S2). mRBD-SpyCatcher and MsDPS2-SpyTag were complexed in the ratio 1:3, the conjugation was confirmed by SDS-PAGE, and the nanoparticulate conjugate was purified by SEC (Figure 1l). The SEC purified nanoparticulate mRBD bound ACE2 and CR3022 with high k_on_ (>10^6^ M^-1^s^-1^) and negligible k_off_, indicating formation of a functional MsDPS2-mRBD nanoparticle (Figure 1m, Supporting Figure S2).

Further, we also assessed binding with lower immobilization and lower concentration of analyte for purified hCMP-mRBD and mRBD-hCMP. The SPR traces were very similar to those observed with higher immobilization and higher analyte concentrations and still revealed negligible dissociation (Supporting Figure S3).

Thermal tolerance to transient and extended thermal stress is a desirable characteristic for deployment of vaccines in low resource settings in the absence of a cold-chain. hCMP-mRBD protein in solution was observed to retain functionality after 1 hour exposure to temperatures as high as 70 °C (Figure 2a). The lyophilized hCMP-mRBD was also observed to retain functionality to transient ninety-minute thermal stress upto 99 °C (Figure 2b). Further, the protein remained natively folded and at 37 °C retained functionality in solution upto three days, and for at least four weeks in the lyophilized state (Figure 2c, 2d, 2e, 2f). In contrast mRBD-GlyIZ showed substantially decreased ACE2 binding after one hour incubation at temperatures above 40 °C and lost ACE2 binding after lyophilization and resolubilization (Supporting Figure S4a, S4b). It is therefore likely that the GlyIZ derived trimer dissociates when subjected to thermal stress above 40 °C and/or lyophilization and is unable to refold back to its native, trimeric state. In contrast, the covalently linked trimers appear to refold back to the native state with much higher efficiency (Figure 2).

**Figure 2.**
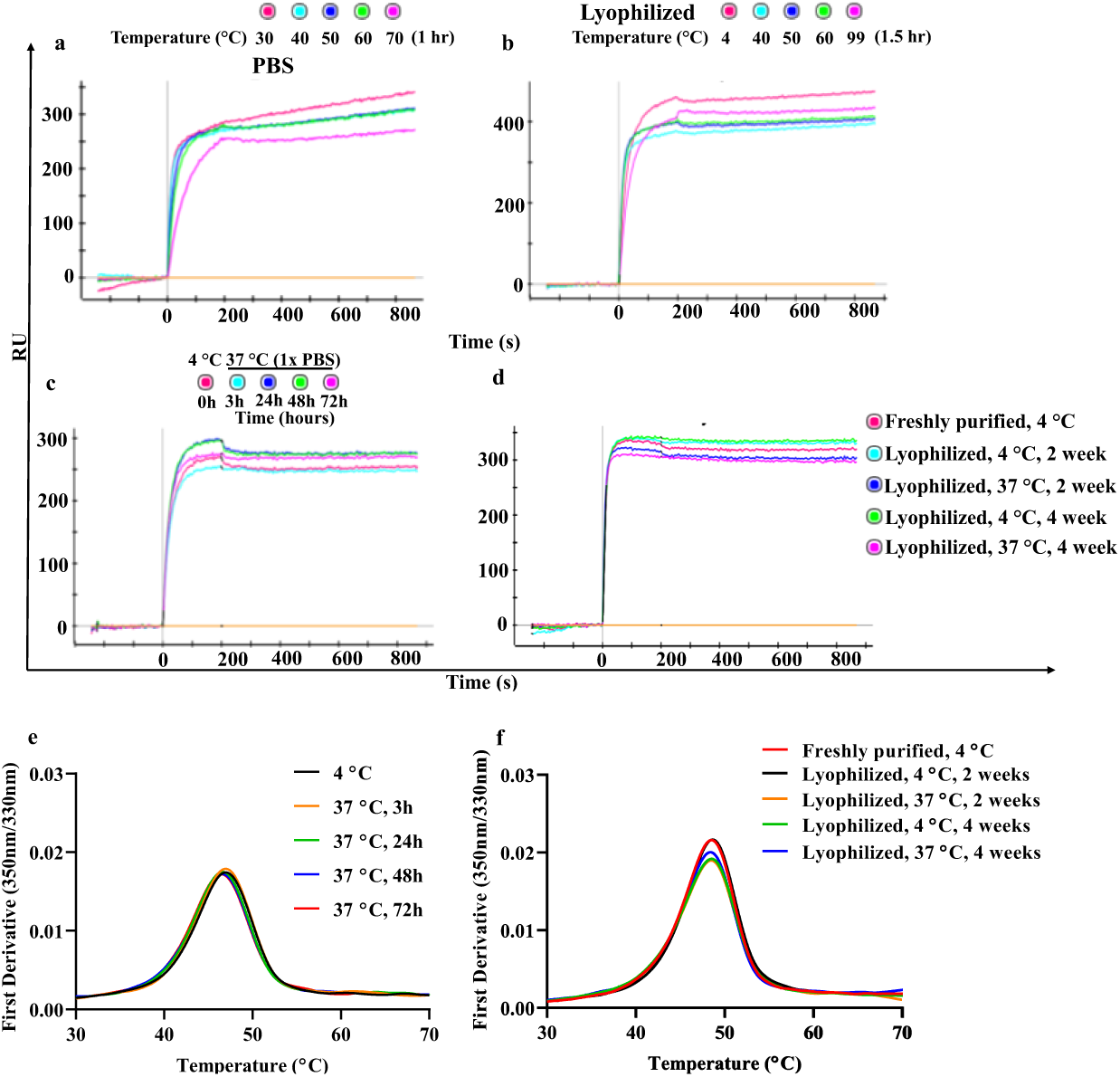
Characterization of trimeric hCMP-mRBD following transient exposure to elevated temperature and extended incubation at 37 °C. **a.** hCMP-mRBD in PBS at a concentration of 0.2 mg/ml was subjected to transient thermal stress for one hour and bindingstudies performed at 100nM. **b.** Lyophilized hCMP-mRBD was subjected to transient thermalstress for 90 minutes followed by reconstitution in water. **c.** hCMP-mRBD (0.2 mg/ml) in solution subjected to 37 °C incubation as a function of time (3-72 hr) **d.** Lyophilized hCMP-mRBD subjected to extended thermal stress at 4 °C and 37 °C for 2 and 4 weeks. 100nM of hCMP-mRBD was used as analyte. **e-f** Equilibrium thermal unfolding monitored by nanoDSF. **e.** hCMP-mRBD (0.2mg/ml) subjected to 37 °C incubation in 1xPBS for upto 72 hours. **f.** nanoDSF of lyophilized hCMP-mRBD incubated for upto 4 weeks at 4 °C and 37 °C. The lyophilized protein was reconstituted in MilliQ grade water prior to thermal melt and SPR binding studies. The binding to ACE2-hFc was performed at 100nM. ACE2-hFc immobilized was 800RU. hCMP-mRBD is thus highly resistant to transient and extended thermal stress.

### Trimeric mRBD elicits high titers of neutralizing antibodies in mice and guinea pigs and protects hamsters from viral challenge

We assessed the immunogenicity of the previously described^37^ monomeric mRBD and trimeric hCMP-mRBD adjuvanted with SWE, an AddaVax™ and MF59 equivalent adjuvant, in BALB/c mice. Animals were immunized intramuscularly at day 0, followed by a boost at day 21^37^. Two weeks post boost, sera were assayed for binding and neutralizing antibodies. Trimeric hCMP-mRBD adjuvanted with SWE elicited 16-fold higher mRBD binding titers compared to monomeric mRBD (Figure 3a). Pseudoviral neutralization titers elicited by trimeric hCMP-mRBD were 45-fold higher (hCMP-mRBD GMT: 31706, mRBD GMT: 707, P *=* 0.008) compared to monomeric mRBD (Figure 3b). We compared the immunogenicity of hCMP-mRBD adjuvanted with AddaVax™ and SWE respectively. The mRBD binding titers and pseudovirual neutralization titers were similar in both adjuvants, confirming their functional equivalence (Supporting Figure S5a, S5b).

**Figure 3.**
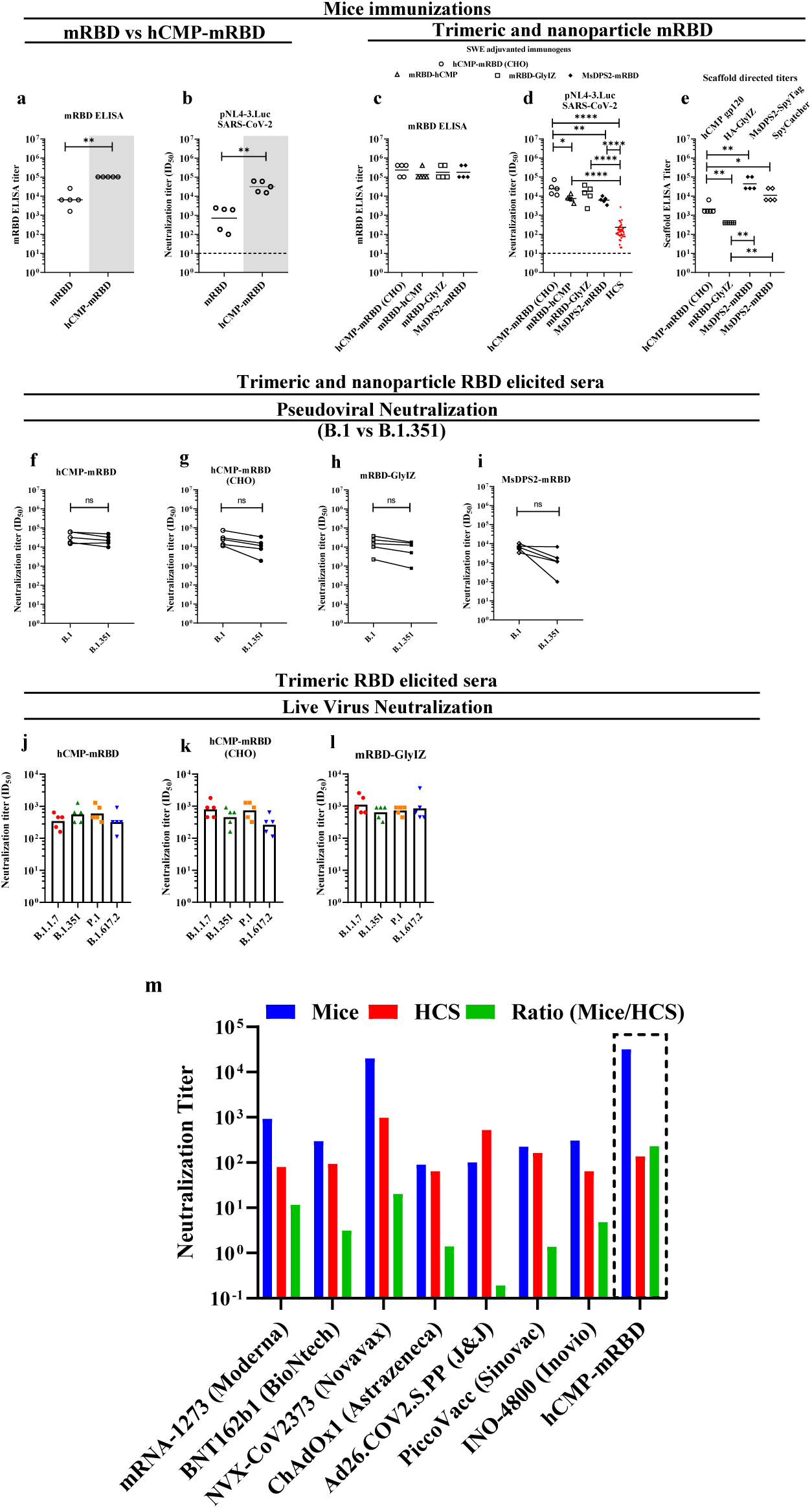
ELISA and pseudovirus neutralization with sera elicited at weeks 0, 3 after two immunizations with SWE adjuvant containing formulations. **a, b.** Immunization with mRBD (white panel) or hCMP-mRBD (gray panel) (n = 5 mice/group) **c-e** Immunizations with mRBD-hCMP, mRBD-GlyIZ or MsDPS2 nanoparticle displaying mRBD (n = 5 mice/group). Pseudoviral neutralization titers utilized pNL4-3.Luc. SARS-CoV-2 D614G Δ19. HCS: Human Convalescent Sera (n = 40). **e.** ELISA binding titer against scaffolds hCMP, GlyIZ trimerization domain, MsDPS2 SpyTag, and SpyCatcher. **f-i.** Pseudoviral neutralization titers against wildtype and pseudovirus with B.1.351 RBD mutations. The paired comparisons were performed utilizing the Wilcoxon Rank-Sum test in f-i. The black solid horizontal lines in each scatter plot represent Geometric Mean Titer (GMT). The pairwise titer comparisons were performed utilizing two-tailed Mann-Whitney test in a-e (* indicates P < 0.05, ** indicates P < 0.01, **** indicates P < 0.0001). **j-l**. Live virus neutralization against B.1.1.7 (Alpha), B.1.351 (Beta), P.1 (Gamma) and B.1.671.2 (Delta). The paired comparisons were performed utilizing ANOVA in j-l, no significant differences were seen. The histograms in each plot represent Geometric Mean Titer (GMT). **m.** Neutralizing antibody titers in mice (blue), in Human Convalescent Sera (HCS) (red) assayed in the identical assay platform, and their relative ratio (green). Values for a number of vaccine candidates being tested in the clinic or provided with emergency use authorizations are shown^58–71^ and corresponding values for hCMP-RBD are boxed.

Next, we assessed the immunogenicity of trimeric, SWE adjuvanted hCMP-RBD derived from different expression platforms, namely CHO and Pichia stable cell lines (Figure 3c, Supporting Figure S6). The binding titers were 12-fold higher in CHO derived hCMP-mRBD compared to hCMP-pRBD (P *=* 0·008) (Figure 3c, Supporting Figure S6). CHO derived hCMP-mRBD (GMT: 24086) elicited high pseudoviral neutralizing titers compared to sera elicited by Pichia expressed hCMP-pRBD which showed negligible neutralization (P = 0·008) (Figure 3d, Supporting Figure S6).

N-terminal trimerized mRBD derived from CHO cells (hCMP-mRBD GMT: 235253) elicited similar mRBD binding titers compared to C-terminal trimerized mRBD in SWE adjuvanted formulations (Figure 3c). The pseudoviral neutralization titer elicited by N-terminal trimerized hCMP-mRBD (GMT: 24086) is 3-fold (P = 0.0317) and 2-fold (P = 0.42) higher compared to C-terminal trimerized mRBD-hCMP (GMT: 7472) and mRBD-GlyIZ (GMT: 12505) respectively (Figure 3d).

MsDPS2-mRBD nanoparticle adjuvanted with SWE, elicited similar mRBD binding antibody titers compared to hCMP-mRBD (GMT: 235253) (Figure 3c) but ∼4-fold (P = 0.008) lower pseudoviral neutralization titers (GMT: 24086 compared to MsDPS2-mRBD, GMT: 6181) (Figure 3d).

The hCMP trimerization domain and nanoparticle scaffolds can also elicit binding antibodies. The binding titers directed towards the Glycosylated IZ were measured by ELISA utilizing influenza HA stem fused to GlyIZ as the immobilized antigen and observed to be the lowest (GMT: 400), 5-fold lower compared to hCMP binding titers (GMT: 2111). The latter were estimated using hCMP V1cyc JRFL gp120 containing the same trimerization domain (Figure 3e)^48^. The MsDPS2-SpyTag and SpyCatcher titers are 111-fold (GMT: 44572, P = 0·0079) and 28-fold higher (GMT: 11143, P = 0.0079) compared to GlyIZ directed titers (Figure 3e) respectively.

We measured the ability of the anti-sera to neutralize pseudovirus containing the RBD mutations present in the isolate B.1.351 (K417N, E484K and N501Y). Sera elicited by hCMP-mRBD, mRBD-hCMP and mRBD-GlyIZ neutralized B.1.351 pseudovirus with 1.4-2.4 fold lower titers compared to Wt pseudovirus (P = 0.05-0.06) (Figure 3f, 3g, 3h). Nanoparticle MsDPS2-mRBD elicited sera neutralized the B.1.351 virus with 5.6-fold lower titers compared to Wt (Figure 3i).

We assessed the immunogenicity of hCMP-mRBD adjuvanted with AddaVax™ in guinea pig immunizations following prime (Wk 0) and, two boosts (Wk 3 and Wk 6). Both binding and pseudoviral neutralization titers were significantly enhanced following the second boost (Supporting Figure S7a, S7b). Trimerization scaffold directed titers in guinea pigs showed only a marginal increase after the second boost (Supporting Figure S7c). Sera collected after the second boost neutralized B.1.351 pseudovirus (GMT: 8252) with 4.3-fold lower titer compared to Wt (GMT: 35693) while corresponding sera from Spike-2P immunized animals showed a 15-fold drop (Supporting Figure S7d, S7e). Unfortunately, sera after the first boost were not available to assay against the B.1.351 pseudovirus. Importantly, both mice and guinea pigs did not elicit any binding antibodies to the L14 linker present in the immunogens.

Pseudoviral neutralization titers correlated well in two independent assay platforms performed with an identical set of sera and with live virus neutralization titers from a CPE based assay (Supporting Figure S8). Additionally, we performed a dose sparing study involving 5 μg hCMP-mRBD adjuvanted with SWE. The mRBD binding titers were observed to be marginally higher compared to the 20 μg dose and pseudoviral neutralization titer were similar. (Supporting Figure S9). Trimeric hCMP-mRBD elicited exceedingly high neutralizing antibodies in mice, compared to Human Convalescent Sera (HCS) titers assayed in the identical assay platform. Additionally, both the mice neutralizing antibody titers and their ratio relative to HCS neutralizing titers compared favorably with corresponding values for vaccine candidates being tested in the clinic or provided with emergency use authorizations (Figure 3m).

As we previously showed, for several COVID-19 vaccines mice titers are predictive of those in humans^37^.

Similar to recent ferret studies involving two other vaccines,^72–74^ microneutralisation assays employing the full length replicative viruses for the four current VOC (alpha, beta, gamma, and delta) were performed with each of the five samples from mice vaccinated with hCMP-mRBD, hCMP-mRBD (CHO) and mRBD-Gly IZ (Figure 3j-3l;, note that the sera used for neutralization studies with full length virus were after three immunizations because all the available sera after two immunizations had been used up in the pseudoviral assays). Negative control (unvaccinated mice sera) and positive control (terminal pooled ferret sera from Marsh et al.^73^) were included for comparison; the assays were performed in duplicates as the volumes of mice sera were limited, however these replicate titers were frequently identical or at most within 2-fold of one another. 50% neutralization titers were calculated for each serum/variant combination on the duplicate values using the method of Spearman and Karber (Figure 3j-3l)^75,76^.

Geometric mean neutralization titers were calculated for each treatment group on log2-transformed data for the alpha, beta, gamma and delta VOC respectively, with titres of 343, 557, 597, and 320 for hCMP-mRBD (Figure 3j); 788, 453, 735, and 260 for hCMP-mRBD (CHO) (Figure 3k); and 1114, 640, 735, and 845 for mRBD-GlyIZ (Figure 3l). Mixed effects ANOVA comparison of the neutralization titers against the four VOC revealed no significant differences in neutralization of any of the VOC for the vaccine formulations used.

To examine efficacy, we carried out a hamster immunization and challenge study. Hamsters were immunized with hCMP-mRBD at Wk 0, 3 and 6. The mRBD binding titers (GMT:18101) and neutralization titers (GMT: 1423) were lower than those observed in guinea pigs and mice (Figure 4a), neutralization titers remained unchanged between the first and second boost. The scaffold directed titers were ∼ 10^3^, consistent with the low sequence identity of hCMP (51%) with the hamster ortholog (Figure 4b). Following immunization, animals were challenged with replicative Wt virus. Two additional groups, namely unimmunized-unchallenged (UC) and unimmunized-virus challenged (VC) animals, acted as controls. Post infection, the immunized animals regained weight and showed markedly lower clinical signs (Figure 4c, 4d), lung viral titers (Figure 4e) and histopathology scores relative to the VC control group (Figure 4f-4j). The tissue sections show clear lung epithelial interstitial spaces and minimal immune cell infiltration in the immunized group compared to virus challenged group.

**Figure 4.**
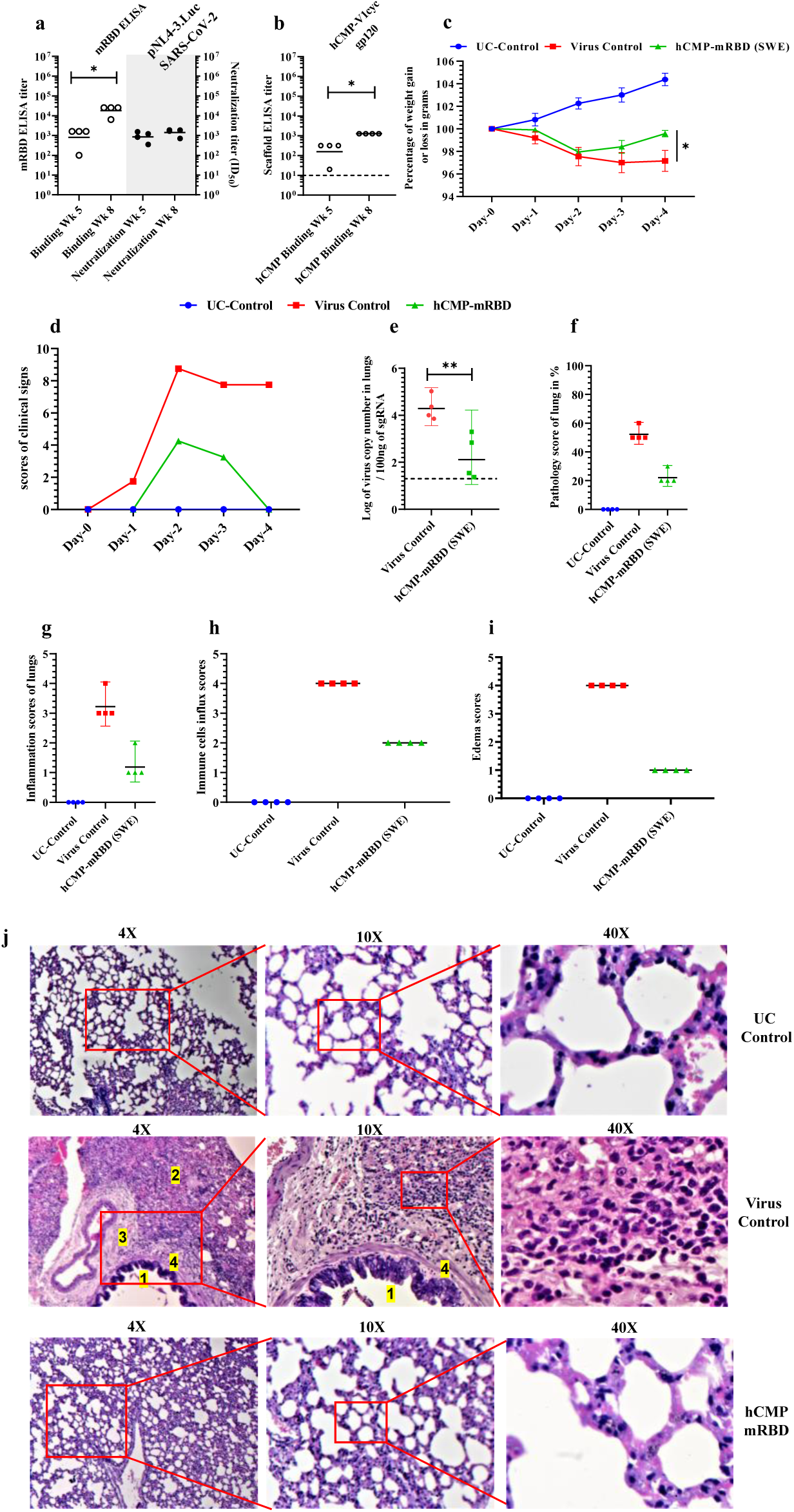
Hamster Immunization and challenge studies with trimeric hCMP-mRBD. **a.** Hamsters (n = 4/group) were immunized at week 0, 3 and 6 with 20 μg of hCMP-mRBD adjuvanted with AddaVax™. **a.** At 14 days post boost, sera were assayed for ELISA binding titer against mRBD and pseudoviral neutralization titer utilizing pNL4-3.Luc. SARS-CoV-2 D614G Δ19. **b.** ELISA binding titer against scaffold hCMP. Post immunization, the hamsters were challenged intranasally with replicative SARS-CoV-2 virus (10^6^ pfu/hamster) and monitored for **c.** Weight change. The pairwise titer comparisons were performed utilizing two-tailed student-t test (* indicates P < 0.05, ** indicates P < 0.01). **d.** Clinical signs **e.** Lung viral titer. Histopathology scores including **f.** Lung pathology score **g.** Inflammation score **h.** Immune cell influx score **i.** Edema score. **j**. Histology of lung sections at varying magnifications (4X, 10X and 40X). Lung pathologies of Unchallenged control (UC), virus challenged and immunized animals. The virus challenged lung histology marked to identify 1. Bronchiolitis and bronchopneumonia, 2. Severe alveolar inflammation (leukocytic alveolitis), alveolar edema and congestion of parenchyma, severe blood hemorrhage and leakage of alveolar sacs, 3. perivascular inflammation and vascular congestion, 4. bronchial infiltration of immune cells with marked edema.

All animals remained healthy after the immunizations. We conclude that hCMP mediated trimerization of mRBD led to elicitation of robust binding and neutralizing antibodies considerably in excess of those seen in convalescent sera, that protected hamsters from high dose, replicative viral challenge.

### Characterization of hCMP-mRBD expressed from permanent cell lines

Stable Chinese hamster ovary (CHO) and HEK293 suspension cell lines expressing the protein were constructed. Purified protein yields were 80-100 mg/liter, similar to those expressed in *Expi293* cells, and SDS-PAGE revealed the presence of disulfide linked trimers (Supporting Figure S10). CHO expressed protein adjuvanted with SWE adjuvant has comparable immunogenicity in mice to transiently expressed protein (Figure 3a, 3b, 3c, 3d).

hCMP-pRBD protein was also expressed in the methylotrophic yeast *P. pastoris* at a purified yield of ∼7mg/liter. As observed previously with monomeric RBD ^37^, the protein was more heterogeneous and formed high molecular weight aggregates unlike mammalian cell expressed proteins (Supporting Figure S6a, Figure 1d and Supporting Figure S10). In mice, formulations with the AddaVax equivalent adjuvant, SWE^77^, elicited low mRBD and negligible neutralization titers after two immunizations (Supporting Figure S6b, S6c).

## Discussion

There are currently multiple COVID-19 vaccines that have been given emergency use approval and others with encouraging Phase I data^66^ are in advanced clinical trials. There remains a need for cheap, efficacious, COVID-19 vaccines that do not require a cold chain and elicit antibodies capable of neutralizing emerging variants of concern (VOC). Despite the extraordinarily rapid pace of vaccine development, there are currently many countries where not even a single dose has been administered. This will prolong the pandemic and promote viral evolution and escape^78^. It has also become clear that minimizing the extent of non-SARS-CoV-2 derived immunogenic sequence in the vaccine is highly desirable. We previously designed a thermotolerant, monomeric, glycan engineered RBD (residues 332-532) that elicited neutralizing antibodies. In the present study we sought to improve the immunogenicity without negatively altering biophysical and antigenic characteristics of the designed immunogen by designing trimeric and nanoparticulate RBDs. Overall, trimeric hCMP-mRBD elicited higher binding and neutralizing antibodies in both mice and guinea pigs compared to monomeric mRBD. Relative to other trimerization domains such as foldon and GCN4 derivatives^32,65^, this forms a homogenous trimer that is stabilized by intermolecular disulfides and hence will not dissociate, even at high dilutions. A fusion of hCMP with HIV-1 gp120 has been extensively tested in guinea pigs, rabbits and non-human primates as an HIV-1 vaccine candidate and showed promising immunogenicity without any apparent adverse effects^48,54,55^. This trimerization sequence has sequence identities with the corresponding ortholog of 81, 91 and 51% in mice, guinea pigs and hamsters, consistent with the low hCMP directed titers in guinea pigs. Thus, hCMP titers in humans are expected to be negligible, given 100% sequence identity with the host protein. Immunization studies in non-human primates with hCMP-mRBD will be shortly initiated to confirm this. In addition, fusion of the short disulfide forming stretch to the other trimerization domains is being carried out to examine the modularity of this motif. Like our previously described monomeric mRBD, hCMP-mRBD shows remarkable thermotolerance. Lyophilized hCMP-mRBD was stable to extended storage at 37 °C for over four weeks and to transient 90-minute thermal stress of up to 100 °C. In contrast, to the alternative GlyIZ trimerization sequence, the disulfide linked hCMP-mRBD was more homogeneous and thermotolerant, demonstrating that the latter feature is not a given for any multimeric RBD formulation. Mice were immunized with various trimeric and nanoparticle displayed RBD constructs (Figure 1) adjuvanted with SWE, a GMP grade adjuvant equivalent to MF59. MF59 has a long safety record of use in humans^47^. Trimerization of mRBD led to elicitation of about 40-fold higher mRBD binding and pseudoviral neutralizing antibodies compared to those elicited by mRBD immunizations in mice. No detectable antibodies were elicited against the short gp120 derived stretch present in the linker of hCMP-mRBD in either mice or guinea pigs. Pseudoviral neutralization titers produced are considerably in excess of those seen in patient derived convalescent sera by factors of ∼25-250 folds in mice and guinea pigs respectively (Figure 3b, 3d, Supporting Figure S7). In mice, the formulation induced higher binding and pseudoviral neutralizing antibodies compared to guinea pigs, presumably owing to various differences in the two host immune systems. Trimeric RBD bound ACE2-hFc receptor and the conformation specific antibody CR3022^79^ tightly, with undetectable dissociation (Figure 1m, Supporting Figure S2, S3). Since neutralization assays differ in their relative sensitivity, in order to compare across studies, it is useful to examine the pseudoviral neutralization titers, relative to a panel of convalescent sera measure in the same assay. Using this criterion, the present titers compare favorably with those of many other formulations that have been tested in the mouse model, including several licensed COVID-19 vaccines. The pseudoviral neutralization titers elicited by the present trimeric RBD compare well with pseudoviral neutralization titers produced by a two dose immunization schedule of nanoparticle displayed RBD adjuvanted with AddaVax (IC_50_ GMT of 5 μg RBD-12GS-I53-50: 20000, corresponding IC_50_ GMT of human convalescent sera: 60, ∼330 fold higher compared to HCS) and the Novavax Spike-2P adjuvanted with Matrix-M1 in mice (CPE_100_ GMT: 20000, CPE_100_ HCS GMT: 983, ∼20 fold higher compared to HCS)^40,66^. Additionally, hCMP-mRBD elicited higher pseudoviral neutralizing titers than a 5 μg mRNA based RBD-foldon trimer vaccine construct BNT162b1 tested in mice (IC_50_ GMT: 753, HCS IC_50_ GMT: 94, ∼8 fold higher compared to HCS)^65^. In contrast to recently described, highly immunogenic multi-component nanoparticle systems^39,40^, the present single component, trimeric RBD might be easier to purify and manufacture and in our hands, nanoparticle display did not confer any significant benefit in immunogenicity over trimerization, whilst the former elicited considerably higher titers of scaffold directed antibodies. However, multicomponent as well as Spy-tagged nanoparticles do have the potential advantage of modularity and being able to display two or more antigens simultaneously. Trimeric RBD elicited sera neutralized B.1.351 pseudovirus with only a relatively small drop in neutralization titer compared to that seen for B.1 virus(Figure 3f-3i). Several prior studies have observed a large drop in neutralization against B.1.351 relative to B.1 For example it was observed^80^ that 50% of convalescent sera showed complete loss of neutralization of B.1.351. In another recent study^81^, a ten-fold drop in neutralization titers against B.1.351 was observed in sera from vaccines immunized with Pfizer-BioNtech and Moderna vaccines. Fortunately, other recent variants such as B.1.1.7 (Alpha) show neutralization titers similar to B.1^82^. Interestingly when compared with guinea pigs immunized with the Spike ectodomain, sera from animals immunized with hCMP-mRBD appear to show a smaller decrease in pseudoviral neutralization titer against B.1.351 though more data are required to confirm this (Supporting Figure S7 d-e). A recent study utilizing E2p nanoparticle display of Spike-2G (K986G, V987G)-ΔHR2 (E1150-Q1208) termed S2GΔHR2 employ two intraperitoneal immunizations led to elicitation of unaltered neutralization titer to the B.1.351 viral strain^83^. While encouraging, such an immunization route is unsuitable for mass vaccinations.

Comparison with live virus neutralisation demonstrates that while the two values are correlated, pseudovirus neutralisation titers are frequently 10-to-100-fold higher, therefore where feasible, it is useful to also employ live, infectious virus when assessing the neutralisation efficacy of vaccine-induced antibodies^72–74^. In the present case, very encouragingly, the trimeric RBD elicited sera were able to neutralize all four current variants of concern equivalently. At the present time, we do not have a definitive explanation for why the sera show lower sensitivity to VOC mutations than sera elicited by current vaccines which employ full length spike as the antigen. There are various possibilities, firstly the VOC have additional mutations outside the RBD including in a major neutralizing epitope in the NTD. It is likely that these mutations affect the neutralizing responses in sera elicited by full length spike. Also, use of the SWE, MF-59 like adjuvant as well as the inherently thermotolerant nature of several of the RBD antigens employed in these studies might have altered the distribution of epitopes targeted, relative to those in the existing vaccines. Finally, the present studies employed mice and hamsters with germlines different from humans. This might also have an influence on the distribution of epitopes targeted. Further studies are required to resolve these issues.

Finally, the present trimeric RBD was safe and efficacious, protecting hamsters from viral challenge. In summary, the present study describes the design of a disulfide linked, highly expressed, homogenous, trimeric RBD immunogen that is stable to long term thermal stress, and induces robust neutralizing antibodies against SARS-CoV-2 including similar neutralization titers against all four current VOC. The availability of permanent cell lines for the immunogen now makes it possible to proceed with further clinical development of this highly immunogenic, thermotolerant and easily producible COVID-19 vaccine candidate.

## Conclusion

We describe a thermotolerant, homogenous, intermolecular disulfide-linked, trimeric RBD that is highly expressed, immunogenic, and elicits sera which neutralize SARS-CoV-2 and its current VOC. This is an excellent candidate for future clinical development and deployment, is easily manufacturable at a large scale, and eliminates the requirement of a cold-chain.

## Methods

### Trimeric RBDs and antibody expression constructs

The present trimeric mRBD construct consists of an N-terminal trimerization domain of human cartilage matrix protein (hCMP) (hCMP residues 298-340) (accession number AAA63904) linked by a 14-residue flexible linker (ASSEGTMMRGELKN) derived from the V1 loop of HIV-1 JR-FL gp120 linked to RBD residues 332-532 (accession number YP_009724390.1) with an engineered glycosylation site (NGS) at N532 fused to an HRV-3C precision protease cleavage site linked to a 10x Histidine tag by a GS linker. The hCMP-mRBD construct reincorporated a glycosylation motif “NIT” at the N-terminal of the mRBD recapitulating the native glycosylation site at N331 in SARS-CoV-2 RBD.

The C-terminal fusion of hCMP trimerization domain was obtained by fusing mRBD (residues 332-532) to hCMP (residues 298-340) by a five-residue linker (GSAGS). This construct is termed mRBD-hCMP. Additionally, the C-terminal fusion of Glycosylated IZ trimerization domain was obtained by fusing mRBD (residues 332-532) to Glycosylated IZ (residues “NGTGRMKQIEDKIENITSKIYNITNEIARIKKLIGNRTAS”) by a five residue linker (GSAGS). This construct is termed mRBD-GlyIZ.

mRBD (residues 332-532) was fused to SpyCatcher (residues 440-549) and the construct was termed mRBD-SpyCatcher.

These constructs were fused to a precision protease (HRV-3C) cleavage site linked to a 10x Histidine tag by a GS linker. These constructs were cloned into the mammalian expression vector pcDNA3.4 under control of a CMV promoter and efficient protein secretion was enabled by the tPA secretion signal peptide sequence. CR3022 antibody heavy and light chain genes were synthesised and subcloned into pcDNA3.4 vector by Genscript (USA).

### Purification of recombinant proteins expressed in *Expi293F* cells

mRBD, hCMP-mRBD, mRBD-hCMP, mRBD-GlyIZ, mRBD-SpyCatcher, mSpyCatcher protein was purified from transiently transfected Expi293F cells following manufacturer’s guidelines (Gibco, Thermofisher) as described previously^37^. A minimum of three independent batches of purifications were performed for all the constructs.

### Tag removal

HRV-3C precision protease digestion was performed to remove the C-terminal 10xHis tag (Protein: HRV-3C = 50:1). HRV-3C digestion was performed for 16 hrs at 4 °C in PBS (pH 7.4). Ni Sepharose 6 Fast flow resin (GE Healthcare) affinity exclusion chromatography was performed to obtain the tagless protein (containing the tag C-terminal sequence: LEVLFQ). The unbound tagless proteins concentration was determined by absorbance (A_280_) using NanoDrop™2000c with the theoretical molar extinction coefficient calculated using the ProtParam tool (ExPASy).

### MsDPS2-SpyTag nanoparticle, mRBD-SpyCatcher construct purification and complexation

Protein nanoparticles present an attractive platform for antigenic display and immune stimulation by mimicking natural infection ^84^. In this study, we conjugated mammalian purified RBD (mRBD) from SAR-CoV-2 virus to a self-assembling bacterial protein - <u>D</u>NA Binding <u>P</u>rotein from <u>S</u>tarved cells, isolated from *Mycobacterium smegmatis* (MsDPS2) (PDB ID: 2Z90). MsDPS2 is purified from *E. coli* as a dodecameric protein making it an excellent choice of vaccine nanoplatform^46^. To enable flexible conjugation, we utilized the SpyTag/SpyCatcher protein coupling strategy, created by splitting the CnaB2 domain of S*treptococcus pyogenes* protein FbaB ^45^. We designed an Msdps2-SpyTag construct by genetic fusion of the 13 residue SpyTag at the C-terminus of a single subunit of the MsDPS2 through a 15 residue linker. MsDPS2-SpyTag protein was expressed in *E. coli* BL21 cells by an overnight induction with IPTG at 20 °C and purified from the cell supernatant as a soluble dodecameric protein using a Ni-NTA column. Similarly, an mRBD_SpyCatcher construct was made by genetically fusing the SpyCatcher at the C-terminal of the mammalian expressed RBD. mRBD_SpyCatcher, which is a monomeric protein, was purified by Ni-NTA chromatography from culture supernatant of transiently transfected *Expi293F*. Purified mRBD2-SpyCatcher is monomeric as confirmed by SDS-PAGE. MsDPS2-Spytag and mRBD2-SpyCatcher, were mixed in 1:3 molar ratio and the reaction was incubated at 25 °C for 3 hours. Conjugation reaction was checked on 12 % SDS PAGE. The protein complex conjugated with mRBD2-SpyCatcher was purified using Superose-6 10/300 analytical column. SDS-PAGE and SEC confirmed the complexation of MsDPS2-SpyTag with mRBD-SpyCatcher.

### SDS-PAGE analysis, Size exclusion chromatography (SEC) and SEC-MALS

Protein purity was estimated by denaturing PAGE. Samples were denatured in SDS containing sample buffer by boiling in reducing (with β-mercaptoethanol) or non-reducing (without β-mercaptoethanol) conditions.

SEC profiles were obtained in 1xPBS buffer equilibrated analytical gel filtration Superdex-200 10/300GL column (GE healthcare) on an Äkta pure chromatography system. The peak area under the curve (AUC) was determined in the Evaluation platform using the peak integrate tool. For SEC-MALS (multi angle light scattering), a PBS (pH 7.4) buffer equilibrated analytical Superdex-200 10/300GL gel filtration column (GE healthcare) on a SHIMADZU HPLC was utilized to resolve hCMP-mRBD purified protein. Gel filtration resolved protein peaks were subjected to in-line refractive index (WATERS corp.) and MALS (mini DAWN TREOS, Wyatt Technology corp.) detection for molar mass determination. The acquired data from UV, MALS and RI were analysed using ASTRA™ software (Wyatt Technology).

### nanoDSF thermal melt studies

Equilibrium thermal unfolding of hCMP-mRBD (- 10xHis tag) protein, before or after thermal stress was carried out using a nanoDSF (Prometheus NT.48) as described previously^37^. Two independent measurements were carried out in duplicate with 2-4 μM of protein in the temperature range of 15-95 °C at 100% LED power and initial discovery scan counts (350nm) ranging between 5000 and 10000. In all cases, when lyophilized protein was used, it was reconstituted in water, prior to DSF.

### SPR-binding of hCMP-mRBD analyte to immobilized ACE2-hFc/CR3022

hCMP-mRBD protein kinetic binding studies to ACE2-hFc and CR3022 antibody were performed on a ProteOn XPR36 Protein Interaction Array V.3.1 (Bio-Rad). The GLM sensor chip was activated with sulfo-NHS and EDC (Sigma) reaction. Protein G (Sigma) was covalently coupled following activation. ∼3500-4000 RU of Protein G (10 µg/mL) was coupled in 10mM sodium acetate buffer pH 4·5 at a flow rate of 30 µl/min for 300 seconds in desired channels. Finally, 1M ethanolamine was used to quench the excess sulfo-NHS esters. Following quenching, ligand immobilization was carried out at a flow rate of 30 µl/min for 100 seconds. ACE2-hFc or CR3022 were immobilized at ∼800 RU on desired channels excluding a single blank channel that acts as the reference channel. hCMP-mRBD analyte interaction with ligands was monitored by passing over the chip at a flow rate of 30 µl/min for 200 seconds, and the subsequent dissociation phase was monitored for 600 seconds. An empty lane without ligand immobilization was utilized for measuring non-specific binding. Following each kinetic assay, regeneration was carried out with 0.1 M Glycine-HCl (pH 2.7). The ligand immobilization cycle was repeated prior to each kinetic assay. Various concentrations of the hCMP-mRBD (- 10xHis tag) (100 nM, 50 nM, 25 nM, 12.5 nM, 6.25 nM) in 1x PBST were used for binding studies. The kinetic parameters were obtained by fitting the data to a simple 1:1 Langmuir interaction model using Proteon Manager.

### SPR-binding of thermal stress subjected hCMP-mRBD analyte to immobilized ACE2-hFc

Lyophilized protein or protein in 1X PBS (0.2 mg/mL) was subjected to transient thermal incubation at the desired temperature in a thermal cycler for ninety or sixty minutes, respectively. Post thermal incubation, binding response was assessed at 100nM analyte concentration by SPR as mentioned in the previous section.

### Mice and Guinea Pig Immunizations

Immunizations of BALBc mice (n=5/group, female, 3-4 weeks old, ∼16-18 g) and Hartley strain guinea pigs (n=5/group, female, 6-8 weeks old, ∼300 g) were performed with freshly adjuvanted (Sepivac SWE™ (Cat.No. 80748J, Batch no. 200915012131, SEPPIC SA, France) and/or (AddaVax™ (vac-adx-10, InvivoGen, USA))) protein (1:1 v/v Antigen : Adjuvant ratio per animal/dose, 20 µg protein in 50 µl PBS (pH 7.4) and 50 µl Adjuvant). Animals were immunized via the intramuscular route with two doses constituting prime and boost on Day 0 and 21 respectively. Sera were isolated from bleeds drawn prior to prime (day -2), post prime (day 14) and post boost (day 35). All animal studies were approved by the Institutional Animal Ethics committee (IAEC) (RR/IAEC/61-2019, Invivo/GP/084, CAF/ETHICS/798/2020, CAF/ETHICS/799/2020, CAF/ETHICS/799/2020, CAF/ETHICS/799/2020). The animal experiments were conducted in compliance to the ARRIVE guidelines^85^. Mice and guinea pig immunizations were non blinded.

### Hamster experiments

#### Ethics and animals’ husbandry

The animal experimental work plans were reviewed and approved by the Indian Institute of Science, Institute Animals Ethical Committee (IAEC). The experiment was performed according to CPCSEA (The Committee for the Purpose of Control and Supervision of Experiments on Animals) and ARRIVE guidelines ^85^. The required number (n = 4/group) of Syrian golden hamsters (*Mesorectums auratus*) of both sexes (50-60 gm of weight) were procured from the Biogen Laboratory Animal Facility (Bangalore, India). The hamsters were housed and maintained at the Central Animal Facility at IISC, Bangalore, with feed and water ad libitum. and 12hr light and dark cycle.

#### Hamster Immunization protocol

After two-week acclimatization of animals, hamsters (n = 4/group) were randomly grouped, and the immunization protocol initiated with the pre-bleed of animals. Hamsters were immunized with 20 ug of subunit vaccine candidate in 50ul injection volume intramuscularly, with the primary on day 0 and boosts on day 21 and 42. Bleeds were performed two weeks after each immunization. The hamster immunization study was non blinded.

#### Virus Challenge

After completing the immunization schedule, the hamsters were transferred to the virus BSL-3 laboratory at the Centre for Infectious Disease Research, Indian Institute of Science-Bangalore (India) and were kept in individually ventilated cages (IVC), maintained at 23 ± 1 °C and 50 ± 5 % temperature and relative humidity, respectively. After acclimatization of seven days in IVC cages at the virus BSL-3 laboratory, the hamsters were challenged with 10^6^ PFU of SARS-Cov-2 US strain (USA-WA1/2020 obtained from BEI resources) intranasally in 100 μl of DMEM, by sedating/anaesthetizing the hamsters with a xylazine (10mg/kg/body wt.) and ketamine (150g/kg/body wt.) cocktail intraperitoneally. The health of hamsters, body temperatures, body weights, and clinical signs were monitored daily by an expert veterinarian. Clinical sign scoring systems were developed similar to that described earlier with some modifications. In the present experiment considering fourteen clinical signs, we measured average clinical scores on as follows, lethargy (1 point), rough coat (1 point), Sneezing (1 point), mucus discharge from nose or eyes (1 point), Half closed eyes/ watery eyes (1 point), huddling in the corner (1 point), ear laid back (1 point), hunched back (1 point), head tilt (1 point), moderate dyspnoea (2 points), body weight loss: 2-5% (1 point), 5-10% (2-point), 10-20% (3 point), Shaking or shivering (1 point).

On the fourth day, post challenge, all the hamsters were humanely euthanized by an overdose of xylazine through intraperitoneal injection. The left lobe of the lung was harvested and fixed in 4% paraformaldehyde (PFA) for histopathological examination of lungs. The right lobes were frozen at -80°C for determining the virus copy number by qRT-PCR.

### Histopathological Examination

Left lobes of lung, fixed in 4% of paraformaldehyde were processed, embedded in paraffin, and cut into 4 μm correct symbol, and sectioned by microtome for haematoxylin and eosin staining. The lung sections were microscopically examined and evaluated for different pathological scores by a veterinary immunologist. Four different histopathological scores were assigned as follows 1. Percent of infected part of lung tissues considering the consolidation of lung; 2. Lung inflammation scores, considering the severity of alveolar and bronchial inflammation; 3. Immune cell influx score, considering the infiltration of lung tissue with the numbers of neutrophils, macrophages and lymphocytes; 4. Edema score, considering the alveolar and perivascular edema. The scores and parameters were graded as absent (0), minimal (1), mild (2), moderate (3), or severe (4)^86^.

### RNA Extractions and qRT-PCR to quantitate sub genomic viral RNA in lungs

Three-time freeze-thawed right lower lobe from the lung of each hamster was homogenized in 1ml of RNAiso Plus Reagent (Takara) and total RNA was isolated as per the manufacturer’s protocol using chloroform and isopropanol reagents. The quantity and quality (260/280 ratios) of RNA extracted was measured by Nanodrop. The extracted RNA was further diluted to 27 ng/μl in nuclease free water. The viral sub genomic RNA copy number was quantified by using 100ng of RNA/well for 10 μl of reaction mixture using AgPath-ID™ One-Step RT-PCR kit (AM1005, Applied Biosystems). The following primers and probes were used 2019-nCoV_N1-Fwd- 5’GACCCCAAAATCAGCGAAAT3’; 2019-nCoV_N1-Rev- 5’TCTGGTTACTGCCAGTTGAATCTG3’; 2019-nCoV_N1 Probe (6-FAM / BHQ-1) ACCCCGCATTACGTTTGGTGGACC (Sigma Aldrich) for amplifying RNA from the SARS CoV-2 N-1 gene. The sub genomic virus copy number per 100ng of RNA was estimated by generating a standard curve from a known number of pfu of the virus.

### ELISA- serum binding antibody end point titers

Desired antigens were coated (4 µg/mL, 50 µL/well, 1xPBS) on 96 well plates for two hours and incubated on a MixMate thermomixer (Eppendorf, USA) at 25 °C under constant shaking (300 rpm). Antigen immobilization was assessed by coating ACE2-hFc protein. Coated wells were washed with PBST (200µl/well) four times, and blocked using blocking solution (100 µL, 3% skimmed milk in 1xPBST) and then incubated at 25 °C for one hour, 300 rpm. Post blocking, antisera were diluted four-folds serially, starting 1:100 and incubated at 25 °C for 1 hour, 300 rpm. Post sera binding, three washes were performed (200 µL of 1xPBST/well). Following this, anti-Guinea Pig IgG secondary antibody (ALP conjugated, Rabbit origin) (diluted 1:5000 in blocking buffer) (50 µL/well) was added and incubated at 25 °C for 1 hour, 300 rpm (Sigma-Aldrich). Post incubation, four washes were performed (200 µL of 1xPBST/well) and incubated with pNPP liquid substrate (50 µL/well) (pNPP, Sigma-Aldrich) at 37 °C for 30 minutes, 300 rpm. Finally, the chromogenic signal was measured at 405 nm. The highest serum dilution possessing signal above cut-off (0.2 O.D. at 405 nm) was considered as the endpoint titer for ELISA.

### Convalescent patient sera samples

Convalescent patient sera were drawn (n = 40) and assayed for pseudoviral neutralization as described in the following pseudovirus neutralization section. The ethics approval of human clinical samples were approved by Institute Human Ethical Committee (Approval No: **CSIR-IGIB/IHEC/2020-21/01**). Patients informed consent was obtained for obtaining the sera following The Code of the Ethics of the World Medical Association (Declaration of Helsinki).

### Production of Pseudotyped SARS-CoV-2 and pseudovirus neutralization assay

Pseudo viral neutralization assays were performed with SARS-CoV-2 pseudo virus harbouring reporter NanoLuc luciferase gene. Briefly, HEK293T cells were transiently transfected with plasmid DNA pHIV-1 NL4·3Δenv-Luc and Spike-Δ19-D614G by using Profection mammalian transfection kit (Promega Inc) following the instructions in the kit manual. Post 48 hours, the pseudovirus containing culture supernatant was centrifuged for 10 mins at 600 xg followed by filtration via 0.22 μm filters, and stored at -80 ^°^C until further use. 293T-hACE-2 (BEI resources, NIH, Catalog No. NR-52511) or Vero/TMPRSS2 (JCRB cell bank, JCRB #1818) cells expressing the ACE2 or ACE and TMPRSS2 receptors respectively were cultured in DMEM (Gibco) supplemented with 5 % FBS (Fetal Bovine Serum), penicillin-streptomycin (100 U/mL). Patient derived convalescent sera (n = 40) were tested for neutralization in both 293T-ACE-2 and Vero/TMPRSS2 cells whereas animal sera were tested only in Vero/TMPRSS2 cells. Neutralization assays were done in two replicates by using heat-inactivated animal serum or human COVID-19 convalescent serum (HCS). The pseudovirus (PV) was incubated with serially diluted sera in a total volume of 100 μL for 1 hour at 37 ^°^C. The cells (Vero/TMPRSS2 or 293T-hACE2) were then trypsinised and 1×10^4^ cells/well were added to make up the final volume of 200uL/well. The plates were further incubated for 48 hours in humidified incubator at 37 ^°^C with 5% CO_2_. After 48 hours of incubation, 140 μL supernatant was removed and 50 μL Bright-Glo luciferase substrate (Promega Inc.) was added. After 2-3 minute incubation, 80 μL lysate was transferred to white plates and luminescence was measured by using Cytation-5 multi-mode reader (BioTech Inc.) The luciferase activity measured as Relative luminescence units (RLU) from SARS-CoV-2 pseudovirus in the absence of sera was used as reference for normalizing the RLUs of wells containing sera. Pseudovirus neutralization titers (ID_50_) were determined as the serum dilution at which infectivity was blocked by 50%. The three RBD mutations (K417N, E484K, N501Y) were introduced into the parental Spike-Δ19-D614G clone using overlap PCR and Gibson recombination. The assembled full-length Spike containing the B.1.351 RBD mutations was cloned in pcDNA3.4 vector and was confirmed by sequencing and used to generate the corresponding pseudovirus as described above. The investigators performing the neutralization assays were blinded to the group identities.

### Growth of current SARS-CoV-2 VOC stocks and live virus neutralization assays

Stocks of the four SARS-CoV-2 variants of concern (VOC), viz. alpha (hCoV-19/Australia/VIC17990/2020, passage 2), beta (501Y.V2.HV001, passage 4), gamma (hCoV-19/Japan/TY7-503/2021, passage 3) and delta (hCoV-19/Australia/ VIC18440/2021, passage 2), were propagated and titrated in Vero E6 cells (CCL81; American Type Culture Collection (ATCC), Manassas, VA, USA) prior to use. Briefly, VeroE6 cells were grown in 150cm^2^ flasks in Dulbecco’s Modified Eagle Medium (DMEM) containing 10% heat-inactivated foetal bovine serum (FBS), 10mM HEPES, 100U/mL penicillin, 100μg/mL streptomycin, and 250ng/mL amphotericin B (all components from ThermoFisher Scientific, Scoresby, VIC, Australia) until 60-80% confluent. The received SARS-CoV-2 isolates were diluted in DMEM containing 10mM HEPES, 100U/mL penicillin, 100μg/mL streptomycin, and 250ng/mL amphotericin B, but no FBS (DMEM-D). Cells were inoculated with 4mL diluted virus and were incubated for 30min at 37°C/5% CO_2_ before 50mL DMEM containing 2% FBS, 10mM HEPES, 100U/mL penicillin, 100μg/mL streptomycin, and 250ng/mL amphotericin B was added. The flasks were incubated for an additional 48h before supernatant was harvested. Harvested supernatants were clarified at 2000g for 10min and stored in 1mL aliquots at -80°C. ATCC VeroE6 cells were additionally used for virus neutralisation assays (see below).

Serum samples used for live virus neutralisation assays were provided by the Indian Institute of Science and Mynvax Private Limited from the mouse study, and the positive control ferret sera was sourced from a different study with the consent of the sponsor (Coalition for Epidemic Preparedness Innovations) and test item provider (University of Oxford). Each serum sample was diluted 1:80 in DMEM-D (see cell culture methods above) in a deep-well plate on a single occasion, followed by a 2-fold serial dilution in medium across the plate up to 1:163,840. The dilution series for each serum sample was dispensed into duplicate rows of a 96-well plate, for a total volume of 50μL per well and duplicate wells per sample dilution. For the serum-containing wells, 50μL virus diluted in medium to contain approximately 100 TCID_50_ (checked by back-titration) was added to each well. The plates were incubated at 37°C/5% CO_2_ for 1h to allow neutralisation complexes to form between the antibodies and the virus. At the end of the incubation, 100μL VeroE6 cells (propagated as outlined above for virus stock generation) were added to each well and the plates returned to the incubator for 4 days. Each well was scored for the presence of viral CPE, readily discernible on Day 4 post-infection, with SN_50_ neutralisation titres calculated using the Spearman-Karber formula^75,76^.

### Statistical Analysis

The P values for ELISA binding titers, neutralization titers, were analysed with a two-tailed Mann-Whitney test using the GraphPad Prism software. The P values for pairwise Wt and B.1.351 pseudovirus neutralization titers were analysed utilizing the Wilcoxon Rank-Sum test. The P value for hamster weight change between virus control and unchallenged groups was analysed by two-tailed student-t test. The correlation coefficients for pseudovirus neutralization 293T-ACE2/VeroE6-TMPRSS2 cell line pseudovirus neutralizations were analysed by Spearman correlation using the GraphPad Prism 8 software (GraphPad v8.4.3). Linear regression model with effects (*lme* function) were undertaken in R 4.0.4 with live virus neutralization titers as dependent variables and antigen and VOC as independent variables^87^ .

## Supporting information

Additional experimental information methods and figures available as single pdf file.

## Ancillary Information

### Supporting information

Additional experimental information methods and figures available as single pdf file.

### Corresponding Author Information

Rajesh P. Ringe, Raghavan Varadarajan.

## Author Contributions

R.V., S.K.M., conceptualized the work, designed the studies. S.P., R.V., R.S.R., planned the animal studies. R.S., constructed the stable cell lines. R.S., U.R.P., P.R., A.U., N.A., K.K., performed ELISA. S.T. standardized growth and quantitation of replicative virus. R.S.R performed the hamster immunizations and challenge studies. K.K. purified ACE2-hFc. S.K., U.R.P. performed the MsDPS2-mRBD construction characterizations. S.P., N.G., A.U., performed hCMP-mRBD protein expression and characterization. S.K.M. carried out DSF and SPR characterization. S.Ga. cloned the mRBD-hCMP and mRBD-GlyIZ and performed hCMP-pRBD protein expression and characterization. S.K., M.S.K., S.K.M., performed SEC-MALS. P.K., cloned the hCMP-mRBD gene. D.C. cloned the Spike with beta VOC RBD mutations. M.B. performed the animal immunizations. I.P., S.D. provided the EM data and analysis. S.M., S.B., provided CPE neutralizing antibody assay data. R.P.R., S.Ku. designed, optimized, executed the pseudoviral neutralization assays, and analyzed the data. R.P.R. was also involved across all the stages of the work. S.S, A.T., S.J., R.P. provided convalescent human serum samples. P.J.v.V., S.R., S.Go., A.J.M., N.B.S., S.S.V. designed, optimized, executed the live virus neutralization assays against the four VOC, and analyzed the data. S.K.M, R.V. wrote the manuscript with contributions from each author.

## Data Availability

All the data are in the manuscript. Additional requests for data will be approved upon reasonable request and upon an approval of signed data access agreement.

## Abbreviations

ACE2: angiotensin-converting enzyme 2
COVID-19: Coronavirus disease 2019
GlyIZ: glycosylated IZ trimerization domain
hCMP: human cartilage matrix protein trimerization domain
HCS: human convalescent sera
Msdps2: *Mycobacterium smegmatis* second DNA binding protein
NTD: N-terminal domain
RBD: Receptor Binding Domain
SARS-CoV-2: severe acute respiratory syndrome coronavirus 2
SD1: short domain 1
SD-2: short domain 2
SEC-MALS: Size Exclusion Chromatography Multi Angle Light Scattering
SPR: Surface Plasmon Resonance
SWE: squalene-in-water emulsion
VOC: variant(s) of concern
Wk: Week

## Acknowledgements

We thank Dr. Neil King for kindly providing the ACE2-hFc fusion protein, and Drs. Lynda Stuart, and Dr. Harry Kleanthous of the Bill and Melinda Gates foundation for helpful discussions. We thank Dr. Barney Graham for kindly providing the Spike-2P construct, and Dr. John Moore for providing SARS-CoV-2 full-length spike and pHIV-1 NL4.3-Δenv-Luc construct. The following reagent NR-52281, SARS-Related Coronavirus 2, Isolate USA-WA1/2020 was deposited by the Centers for Disease Control and Prevention and obtained through BEI Resources, NIAID, NIH: SARS-Related Coronavirus 2, Isolate USA-WA1/2020. We thank BEI Resources for providing 293T-ACE2 cells submitted by Dr. Jesse Bloom. S.S.V. would like to thank the following collaborators for the SARS-CoV-2 VOC used in the live virus neutralisation assays: Victorian Infectious Diseases Reference Laboratory (especially Dr Julian Druce) for hCoV-19/Australia/VIC17990/2020 (alpha VOC) and hCoV-19/Australia/ VIC18440/2021 (delta VOC) isolates; Professors Tulio de Oliveira and Alex Sigal in South Africa for 501Y.V2.HV001 (beta VOC) isolate; Japan’s National Institute of Infectious Diseases (especially Dr Mutsuyo Takayama-Ito) for hCoV-19/Japan/TY7-503/2021 (gamma VOC) isolate; and colleagues at the CSIRO’s Australian Centre for Disease Preparedness (especially Kristen McAuley, Dr Kim Blasdell and Dr Mary Tachedjian for the VOC’s import and Next-Generation Sequencing). S.S.V. is also grateful to the Coalition for Epidemic Preparedness Innovations (especially Dr Amy Shurtleff) and the University of Oxford (especially Professor Dame Sarah Gilbert) for access to positive control ferret sera from their study (CSIRO ACDP Animal Ethics Committee Approval Reference: AEC 2004). This work was funded by a grant from the Bill and Melinda Gates Foundation (INV-005948) and Govt. of India PM CARES fund (SP/OPSA-20-0004) to R.V., and by a major grant from Australia’s Department of Finance to CSIRO (P.I.: S.S.V). We also acknowledge funding for infrastructural support from the following programs of the Government of India: DST-FIST, UGC Center for Advanced Study, MHRD-FAST, the DBT-IISc Partnership Program, and of a JC Bose Fellowship from DST to R.V. S.S.V. is grateful for additional funding from the CSIRO’s Precision Health and Responsible Innovation Future Science Platforms, and the National Health and Medical Research Council. Mynvax acknowledges funding support from IISc CSR grant for COVID19 vaccine work. S.K.M, K.K. acknowledge the support of the MHRD-IISc doctoral fellowship. P.K. gratefully acknowledges financial support from the DBT-RA Program in Biotechnology and Life Sciences. S.D. acknowledges the support of DBT Cryo-EM facility (BT/INF/22/SP22844/2017SM). D.C. acknowledges the support of CSIR doctoral fellowship. S.K. and M.S.K. acknowledge the support of DBT doctoral fellowship. S.M. and S.B. acknowledge the intramural funding received from THSTI under Translational Research Program grant. R.P. acknowledges the support of CSIR_IGIB grant (MLP-2005) and Fondation Botnar (CLP-0031). R.P.R. acknowledges the support of SERB grant (IPA/2020/000168). The content is solely the responsibility of the authors and does not necessarily represent the official views of the funding institutions.

## Table of Contents (TOC)/Abstract Graphic

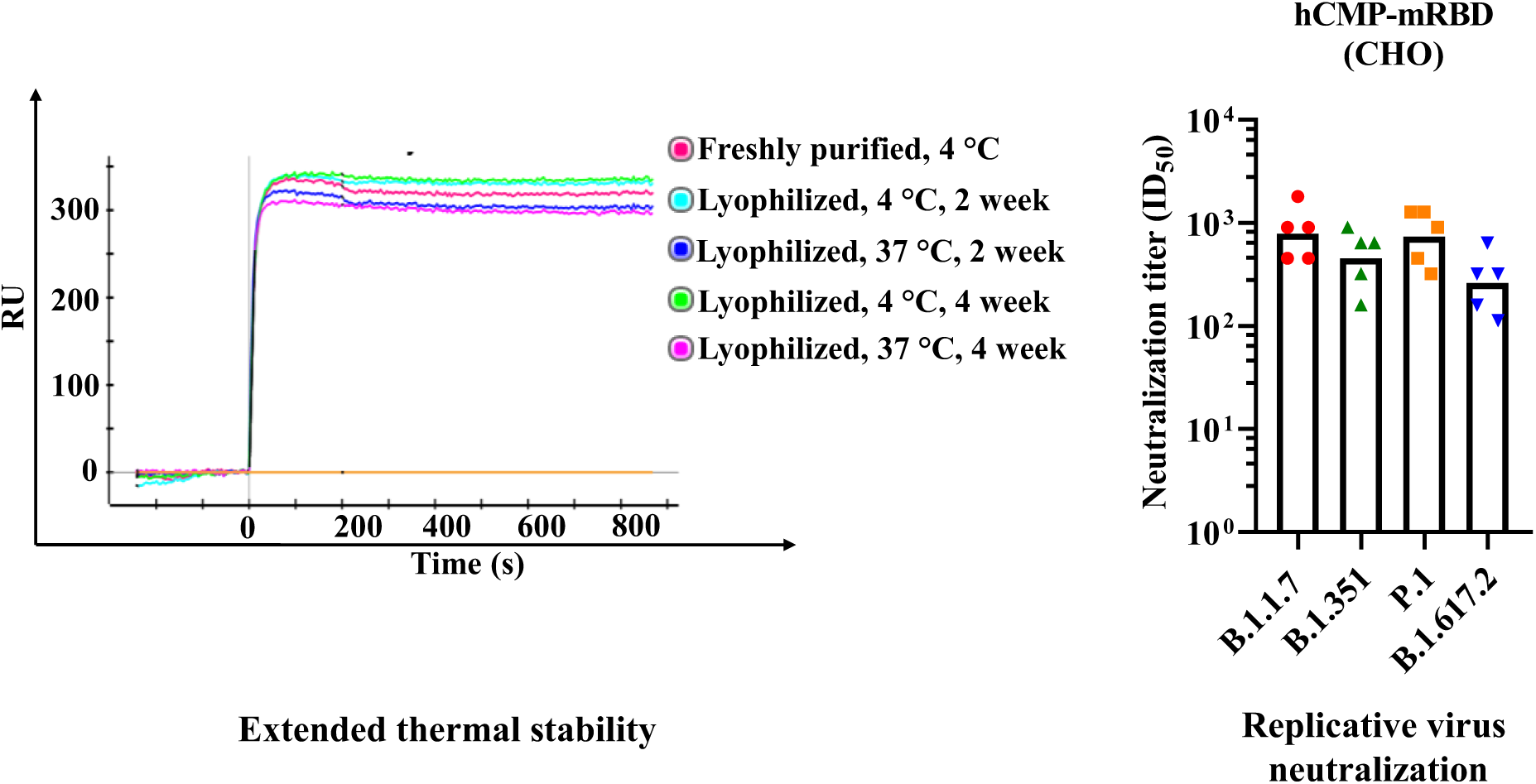

