## Additional experimental information methods and figures available as single pdf file. for "Immunogenicity and protective efficacy of a highly thermotolerant, trimeric SARS-CoV-2 receptor binding domain derivative"

#### Expression and Purification of hCMP-pRBD

The construct sequence was codon-optimized for expression in *Pichia Pastoris* and cloned into the vector pPICZαA containing a MATalpha signal sequence for efficient secretion. The resulting clone was named hCMP-pRBD. The hCMP-pRBD plasmid was linearized with *Pme*I enzyme (NEB, R0560) prior to transformation.

10 µg of linearized plasmid was used for transformation into *Pichia pastoris* X-33 strain by electroporation as described in the user manual for Pichia expression by Thermo Fisher Scientific. The transformants were selected by plating on YPDS (YPD Sorbitol) plates with 100 µg/ml and 1mg/ml Zeocin (Thermo Fisher Scientific, R25005) and incubating the plates at 30 $^{\circ}$C for upto 3 days.

25 colonies from the YPDS plate with 1 mg/ml Zeocin were picked and screened for expression by inducing with 1 % methanol every 24 hrs. Culture tubes (15 ml) with 1ml BMMY media (pH 6.0) each were used for inducing the cultures for upto 120 hrs at 30 $^{\circ}$C and 250 rpm. The expression levels were checked using a dot blot analysis with Anti-his tag antibodies conjugated with HRP enzyme. The colony showing the highest expression level was then chosen for large scale expression. The large scale culture was grown in 2l baffled shake flasks with 350 ml volume of culture. The expression levels were monitored every 24 hrs using Sandwich-ELISA.

The culture was harvested by centrifugation at 12000g, and the supernatant was filtered through a 0.45 micron filter. The supernatant was then incubated with Ni Sepharose 6 Fast flow resin (GE Healthcare) for 2 hrs. The beads were washed with 50 column volumes of 1X PBS pH 7.4 supplemented with 20 mM Imidazole. The His tagged protein was then eluted using 1X PBS pH 7.4 supplemented with 300 mM Imidazole. The eluted fractions were assessed for purity on a 12 % SDS-PAGE. The appropriate fractions were then pooled and dialyzed against 1X PBS to remove Imidazole.

**Cell lines, media and growth conditions for polyclonal stable cell lines**

Flp-In™-293 (Thermo Fisher Scientific, Cat# R75007, Lot# 2220695) as well as Flp-In™-CHO (Thermo Fisher Scientific, Cat# R75807, Lot # 2127131) adherent cells were used for making COVID-19 antigen hCMP-mRBD-HRV-Tg (a stop codon after 'Q' of HRV3C site LEVLFQGP) polyclonal stable cell line. The cell line encoded hCMP-mRBD sequence is thus identical to that obtained after tag removal following HRV3C protease cleavage of protein produced by transient transfection. These engineered cells harbour a single Flp-In™ target site from vector 'pFRT/lacZeo' which confers Zeocin resistance. We first engineered COVID-19 antigen expressing recombinant cells using these adherent cells and then adapted them to suspension conditions for protein production.

**Adherent cell culture**

Both of the above adherent cells were cultured either in T25 or T75 EasYFlask, with a TC surface, filter cap (Thermofisher Scientific Cat# 156367 and 156499) in a moist 8 % $CO_2$ incubator at 37 °C.

The adherent Flp-In™-293 cells were grown in DMEM, high glucose media (Thermo Fisher Scientific Catalog #: 11965118) supplemented with 10 % Fetal Bovine Serum (FBS), qualified Brazil (Thermo Fisher Scientific Cat# 10270106), 100 U/ml Penicillin Streptomycin (Thermo Scientific Cat#15140122) and 100 μg/ml Zeocin™ Selection Reagent (Thermofisher Scientific Cat# R25001).

The adherent Flp-In™-CHO cells were grown in Ham's F-12 Nutrient Mix media (Thermo Fisher Scientific Catalog #: Cat # 11765054) supplemented with 10% FBS, 100 U/ml Penicillin-Streptomycin and 100 μg/ml Zeocin™ Selection Reagent.

**Plasmid and vector**

The Flp-In™ T-REx™ core kit containing pOG44 (Flp recombinase expressing plasmid) and pcDNA5/FRT/TO (donor plasmid for gene of interest) was purchased from Invitrogen USA (Cat # K650001).

The gene of interest 'hCMP-mRBD-HRV-Tg' was PCR amplified from hCMP-mRBD pCMV1 vector using *Hind*III site containing forward primer (5'—TATATAAGCTTCTGCAGTCACCGTCCTTAGATC—3') and *Xho*I site containing reverse primer (5'—TATATCTCGAGTCACTGGAACAGCACCTCCAGGGAGCC—3').

The amplified PCR product was digested with *Hind*III and *Xho*I and subcloned into pcDNA5/FRT/TO restricted with the above two enzymes. The clone was confirmed by sequencing.

**Generation of adherent polyclonal Flp-In stable lines**

T25 flasks (5 ml media) having either adherent Flp-In™-293 or Flp-In™-CHO cells (~80 % confluent) were co-transfected with pOG44 (10 µg) and hCMP-mRBD-HRV-Tg-pcDNA5/FRT/TO (5µg) plasmid DNA using 35 µg of Lipofectamine™ 2000 Transfection Reagent (Thermo Fisher Scientific, Cat # 11668030) in serum free media as per the manufacturer instruction for 4 hrs. After 4 hrs, the media was replaced with serum containing media. The cells were incubated for 16 hrs and then trypsinized using 1 ml of 1X-Tryple express enzyme (Thermofisher Scientific, Cat# 12604021) and seeded to a T75 flask containing 25 ml of desired media and incubated for further 24 hrs for FLP recombination. After 24h the media was replaced with fresh media having Hygromycin 100 µg/ ml (Thermofisher Scientific Cat# 10687010) for Flp-In™-293 and 750 µg/ ml for Flp-In™-CHO cells. Hygromycin resistant foci were observed after 3 days of selection. Media containing the desired amount of Hygromycin was changed after every 5 days mentioned above. After 18 days in case of Flp-In™-293 and 14 days in case of Flp-In™-CHO, the recombinant hygromycin resistant cells reached to 100% confluency. The secretion of the protein of interest (hCMP-mRBD-HRV-Tg) was confirmed from cell free media using western blotting with polyclonal Guinea pig sera against the same antigen. The confirmed polyclonal cells were frozen in liquid $N_2$ for long term storage. The T75-flask grown polyclonal cells were adapted for shake flask suspension culture and used for protein production.

**Shake flask suspension cell culture and protein production**

The suspension cells were grown in 125 or 250-ml Nalgene™ single-use PETG Erlenmeyer flasks with plain bottom and vented closure (Thermofisher Scientific Cat# 4115-0125 or 4115-0250) at 125 rpm with moist 8% $CO_2$ incubator at 37°C or as specifically mentioned.

The stable adherent recombinant Flp-In™-293 cells were first trypsinized from the T75 flask and then grown in a suspension flask after adapting them to FreeStyle™ 293 Expression Medium (Thermofisher Scientific Cat# 12338018) supplemented with 2% FBS and 50 µg/ml Hygromycin B for ~6 generations (two passages, doubling time=24h). These ~300 million cells were then seeded to 100 ml serum free FreeStyle™ 293 Expression medium for protein production for 3 days. After 3 days the media was used for protein purification. The ~300 million cells were grown further in 100 ml media for 6 days under identical conditions and used again for protein purification with >95% cell viability.

The stable adherent recombinant Flp-In™-CHO cells were first trypsinized from a T75 flask and then grown in a suspension flask for direct adaptation to PowerCHO™ 2 Serum-free Chemically Defined Medium (Lonza, Cat# 12-771Q) supplemented with 8 mM L-Glutamine (Thermo Fisher Scientific, Cat# 25-030-081) with 50 µg/ml Hygromycin B. First cells were grown for ~8 generations (two passages, doubling time=24h) at 37 °C till ~3 million per ml density. ~300 million cells were then seeded in 100 ml medium for protein production for 3 days at 32°C. After 3 days the media was harvested for protein purification. The ~300 million cells were grown further in 100 ml media for 6 days under identical condition and media used for protein purification with >95% cell viability.

**Tagless protein purification**

The spent media from stable hCMP-mRBD-HRV-Tg-Flp-In™-293 or Flp-In™-CHO grown cells contained the expressed protein. Protein was purified using anion exchange chromatography. 100 ml cell free media was first dialyzed against 30mM Tris-HCl buffer pH 8.4 overnight at 4 °C using cellulose membrane dialysis tubing (10kDa molecular weight cutoff, Sigma, Cat # D9527-100FT). 2mL Q Sepharose$^{Tm}$ Fast Flow beads (GE Healthcare, Cat# 17-0510-01) were equilibrated with 30mM Tris-HCl pH 8.4 and incubated for 1hr at 4°C with the dialyzed sample. Protein elution was performed with a step gradient of 30mM Tris-HCl pH 8.4. containing 20-500mM NaCl. The fractions were analyzed on a 10% SDS-PAGE gel and the pure fractions were pooled and further dialyzed against 1X-PBS buffer pH 7.4, overnight. The pure protein was analyzed on 10% oxidizing as well as reducing SDS PAGE for homogeneity and purity. Size exclusion chromatography utilizing Superose 6 10/300 Increase GL column with 1X PBS as running buffer at a flow rate of 0.5mL/ min on an ÄktaPure (GE) was performed to determine protein aggregation state.

**Negative Staining sample preparation and visualization by Transmission Electron Microscope**

For negative staining electron microscopy, hCMP-RBD protein was purified by affinity chromatography followed by size exclusion chromatography. The protein eluted as trimer. For visualization by the Transmission Electron Microscope, the sample was prepared by a conventional negative staining method. Briefly, 3.5 μl of sample (0.1mg/ml) was added to glow discharged continuous carbon coated copper grid for 1 minute. The extra sample was blotted out. Negative staining was performed using 1% Uranyl Acetate solution for 20 seconds, following which the grid was air-dried. The negatively stained sample was visualized at room temperature using a Tecnai T12 electron microscope equipped with a Tungsten filament operated at 120 kV. Images were recorded using a side-mounted Olympus VELITA (2KX2K) CCD camera at a magnification of 135k (3.5 Å/pixel).

**Reference-free 2D classification using single-particle analysis**

The evaluation of micrographs was done with EMAN 2.1[1]. Around 6600 particles were picked manually and extracted using e2boxer.py in EMAN2.1 software. Reference free 2D classification of different projections of particles were calculated using simple_prime2D of SIMPLE 2.1 software[2].

### Supporting Figures

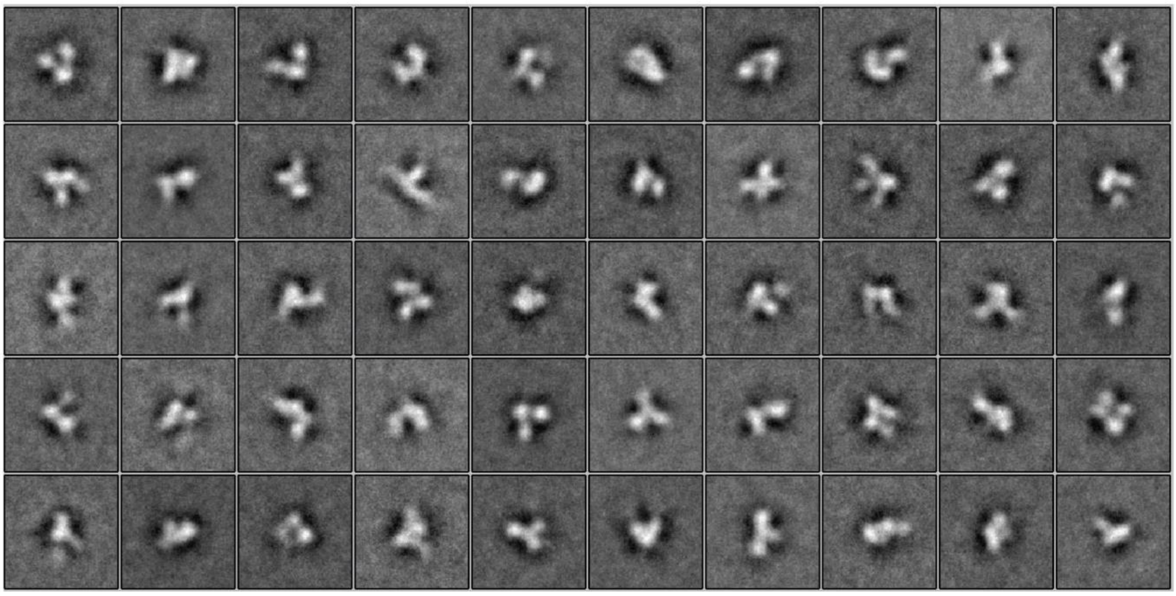

**Supporting Figure S1.**

**Reference free 2D classification calculation of hCMP-mRBD:** The reference free 2D classification calculation using SIMPLE 2.1.

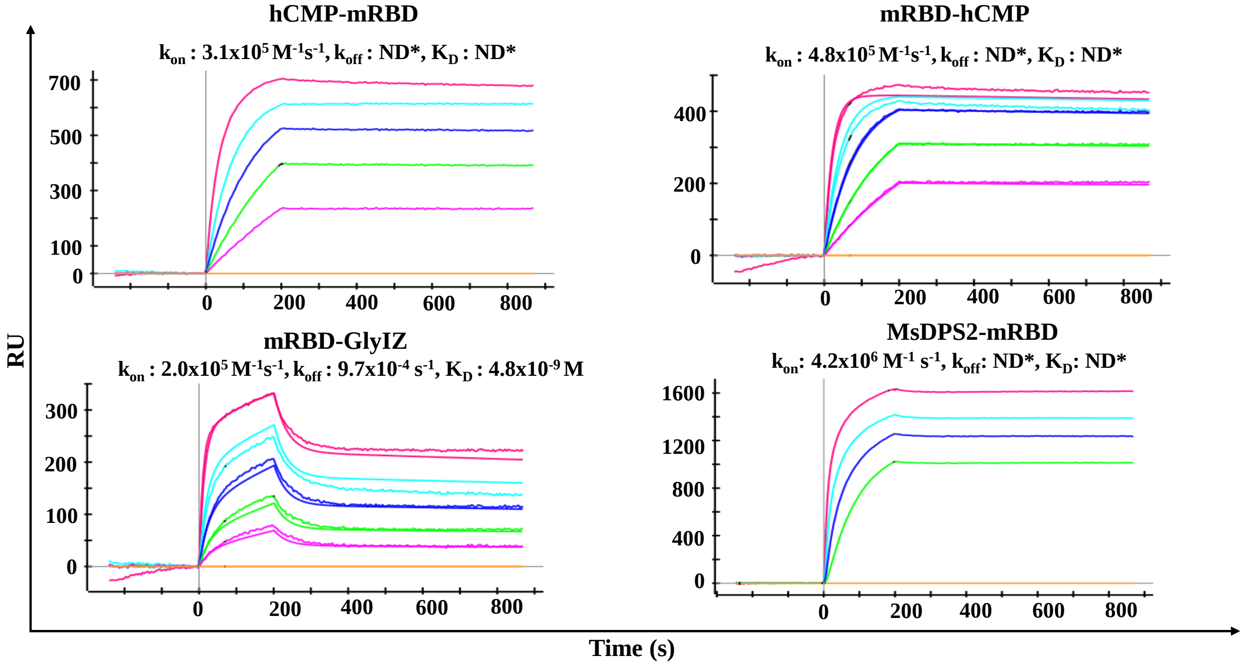


**Supporting Figure S2. SPR binding of trimeric and nanoparticle RBDs to CR3022.** SPR binding studies were performed with hCMP-mRBD, mRBD-hCMP, mRBD-GlyIZ and SEC purified complex MsDPS2-mRBD to CR3022 (Immobilization ~800RU). The curves from highest to lowest correspond to concentrations 100 nM, 50 nM, 25 nM, 12.5 nM and 6.25 nM respectively for hCMP-mRBD, mRBD-hCMP and mRBD-GlyIZ. The curves for MsDPS2-mRBD correspond from highest to lowest to concentrations 10 nM, 5 nM, 2.5 nM and 1.25 nM respectively. ND*- No dissociation.

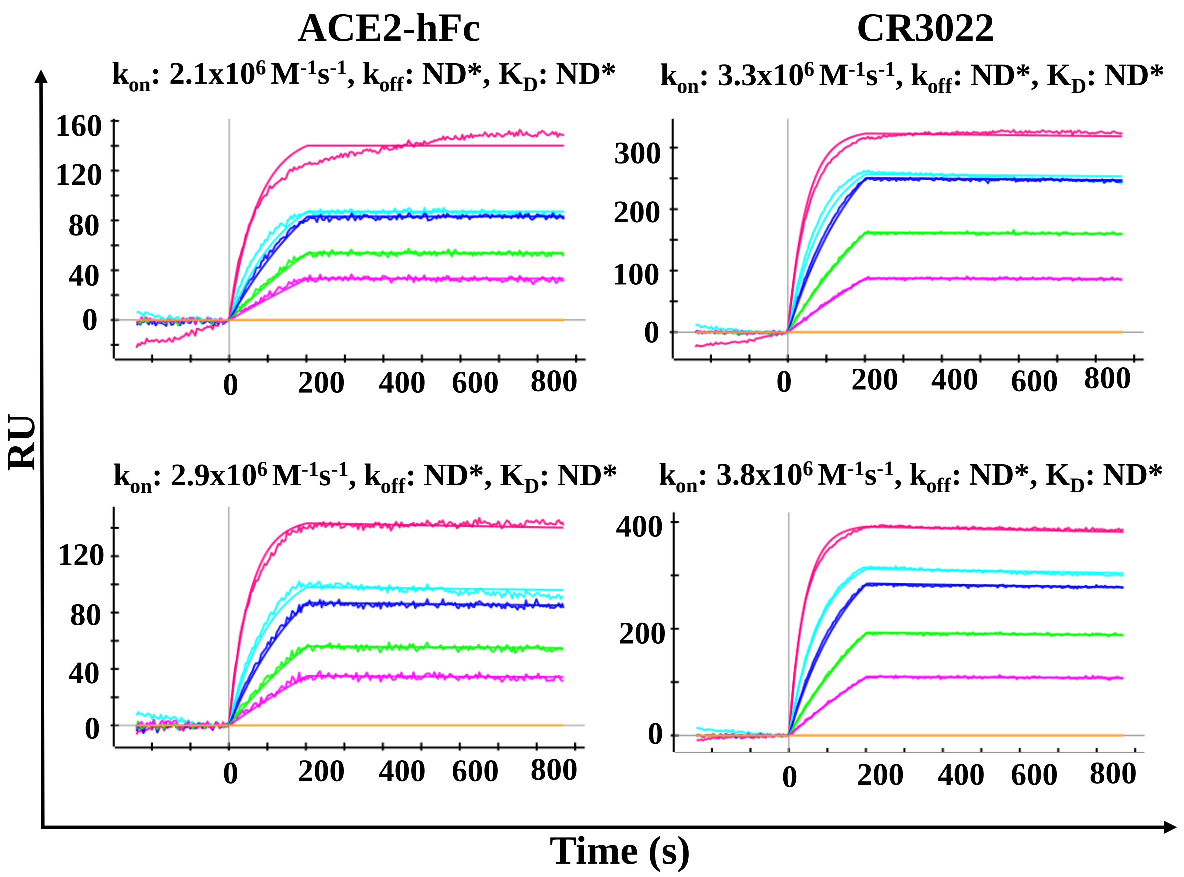


**Supporting Figure S3. SPR binding of trimeric RBDs to ACE2-hFc and CR3022 at low surface immobilization (~400RU) and analyte concentration.** SPR binding studies were performed with hCMP-mRBD, mRBD-hCMP. The curves from highest to lowest correspond to concentrations 6.2 nM, 3.1 nM, 1.5 nM, 0.7 nM and 0.4 nM respectively for hCMP-mRBD and mRBD-hCMP. ND*- No dissociation.

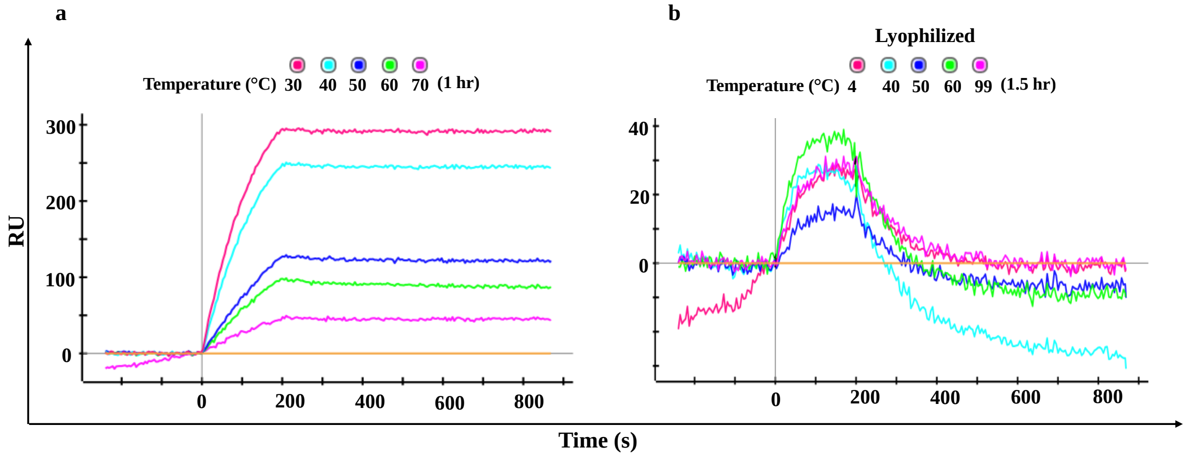


**Supporting Figure S4. Characterization of mRBD-GlyIZ trimeric RBD following transient exposure to elevated temperature. a.** mRBD-GlyIZ in PBS at a concentration of 0.2 mg/ml was subjected to transient thermal stress for one hour and binding studies performed at 100nM. **b.** Lyophilized mRBD-GlyIZ was subjected to transient thermal stress for 90 minutes followed by reconstitution in water. The lyophilized protein was reconstituted in MilliQ grade water prior to thermal melt and SPR binding studies. The binding to ACE2-hFc was performed at 100nM. ACE2-hFc immobilized was 800RU. In solution, mRBD-GlyIZ loses activity upon exposure to temperatures higher than 40 ºC. The molecule also loses activity upon lyophilization and resolubilization.

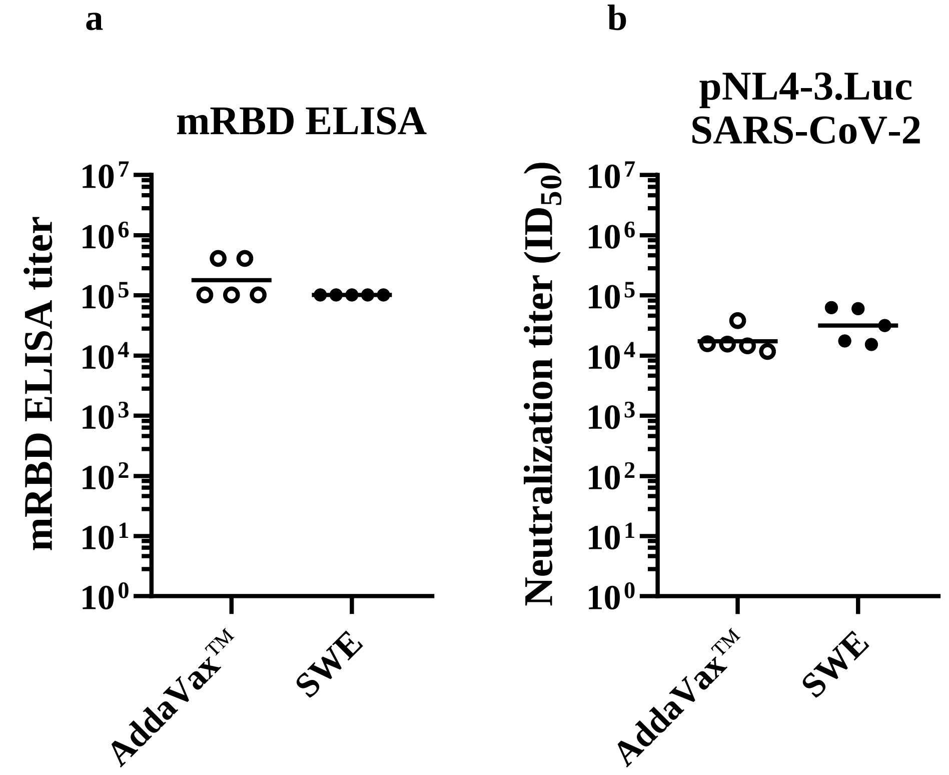


**Supporting Figure S5. hCMP-mRBD adjuvant comparisons.** Mice (n = 5 mice/group) were immunized at week 0 and 3 with 20 μg of hCMP-mRBD adjuvanted with AddaVax™ and SWE. At 14 days post boost, sera were assayed for **a.** ELISA binding titer against mRBD. **b.** Pseudoviral neutralization titer utilizing pNL4-3.Luc. SARS-CoV-2 D614G Δ19.

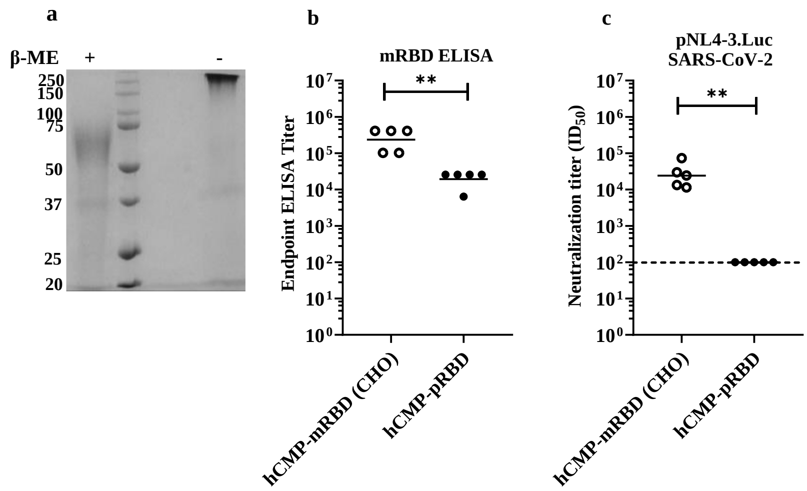


**Supporting Figure S6. Immunogenicity of CHO and Pichia expressed hCMP-RBD. a.** SDS-PAGE of hCMP-pRBD purified from *P.pastoris* under reducing (+) and non-reducing (-) conditions. **b.** Mice (n = 5 mice/group) were immunized at week 0, 3 with 20 μg of hCMP-mRBD (CHO) or hCMP-pRBD adjuvanted with the Addavax equivalent SWE adjuvant. At day 14 post boost, sera were assayed for ELISA binding titers to mRBD. **c.** Pseudoviral neutralization titer utilizing pNL4-3.Luc. SARS-CoV-2 D614G ∆19. The black horizontal lines in each scatter plot represent Geometric mean titer (GMT). The pairwise titer comparisons were performed utilizing two-tailed Mann-Whitney test (** indicates $P < 0.01$).

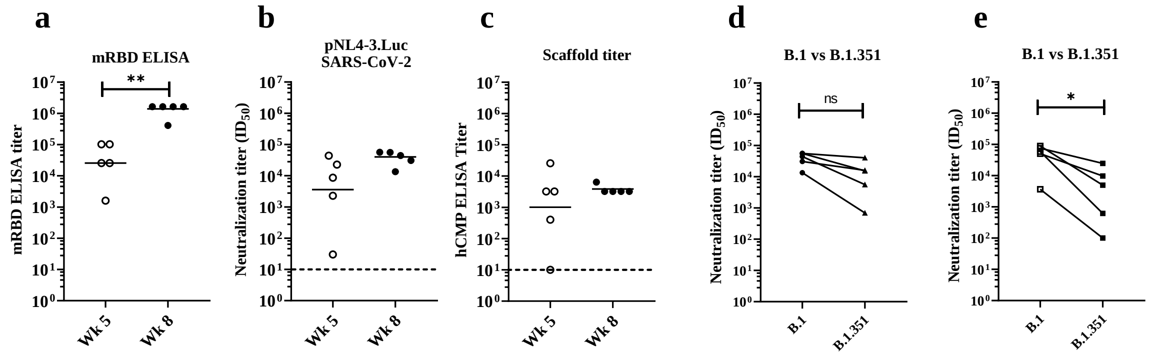


**Supporting Figure S7. Guinea Pig Immunizations.** Guinea pigs (n = 5 guinea pig/group) were immunized at week 0, 3 and 6 with 20 µg of trimeric hCMP-mRBD adjuvanted with AddaVax™. **a.** 14 days post boost, sera were assayed for ELISA binding titer against mRBD. **b.** Pseudoviral neutralization titer utilizing pNL4-3.Luc. SARS-CoV-2 D614G ∆19. **c.** ELISA binding titer against scaffold hCMP. **d-e.** Pseudoviral neutralization titer utilizing the wildtype and South African (B.1.351) derived pseudovirus with sera obtained 14 days post second boost with **d.** hCMP-mRBD. **e.** Spike-2P. The pairwise titer comparisons were performed utilizing two-tailed Mann-Whitney test in A (** indicates P < 0.01) and in D, E were performed utilizing paired two-tailed student-t test (* indicates P < 0·05). The titer with B.1.351 variant reduced by 4-folds in **d.** hCMP-mRBD immunized group and 15-fold in **e.** Spike-2P immunized group compared to Wt.

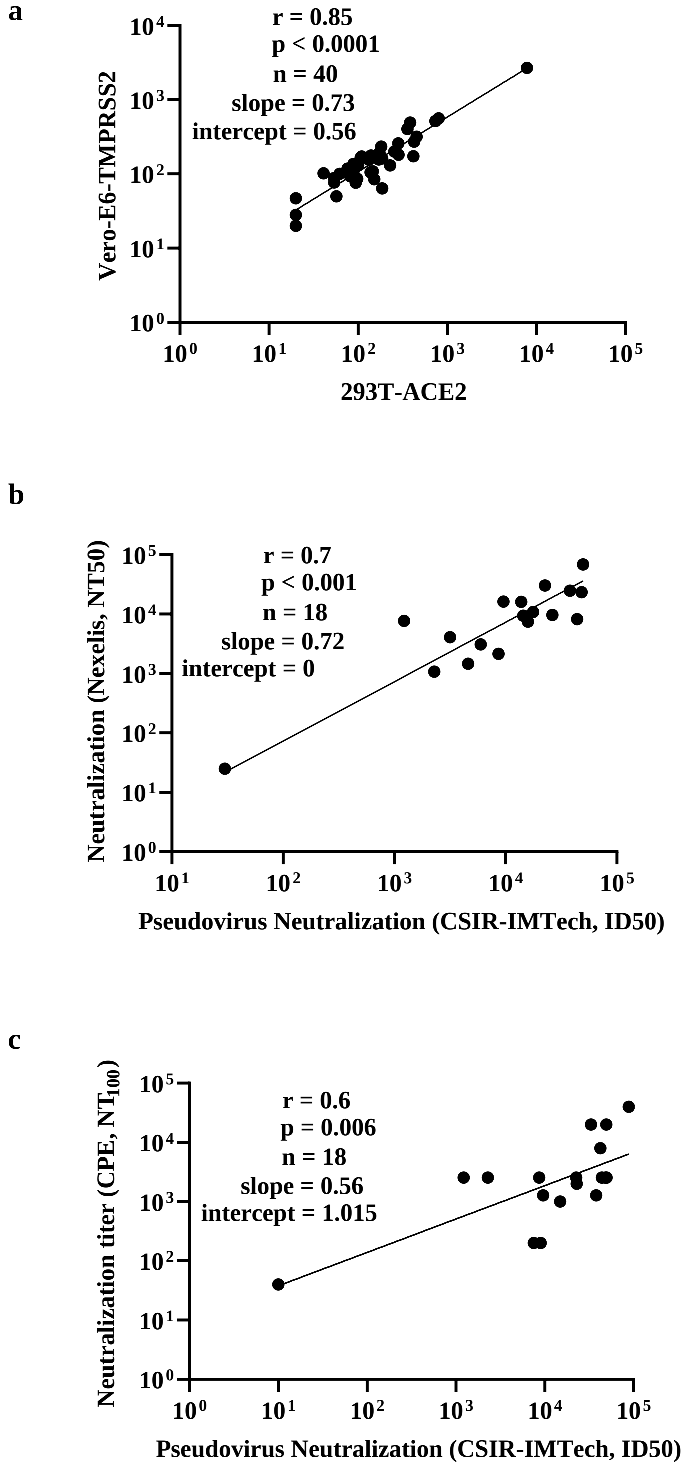


**Supporting Figure S8. Pseudoviral neutralization titer correlations. a.** Human convalescent serum samples (n=40) assayed for pseudoviral neutralization in 293T-ACE2 and Vero-E6-TMPRSS2 cells. Neutralization titers in both cells are very similar. **b.** Identical set of sera (n = 18) assayed for pseudoviral neutralization in Vero-E6-TMPRSS2 (CSIR-IMTech) and at Nexelis (NT50). **c.** Identical set of sera (n = 18) assayed for pseudoviral neutralization in ACE2-293T ($ID_{50}$) (CSIR-IMTech) and CPE based live virus neutralization ($NT_{100}$) at THSTI. Spearman correlation coefficients are shown.

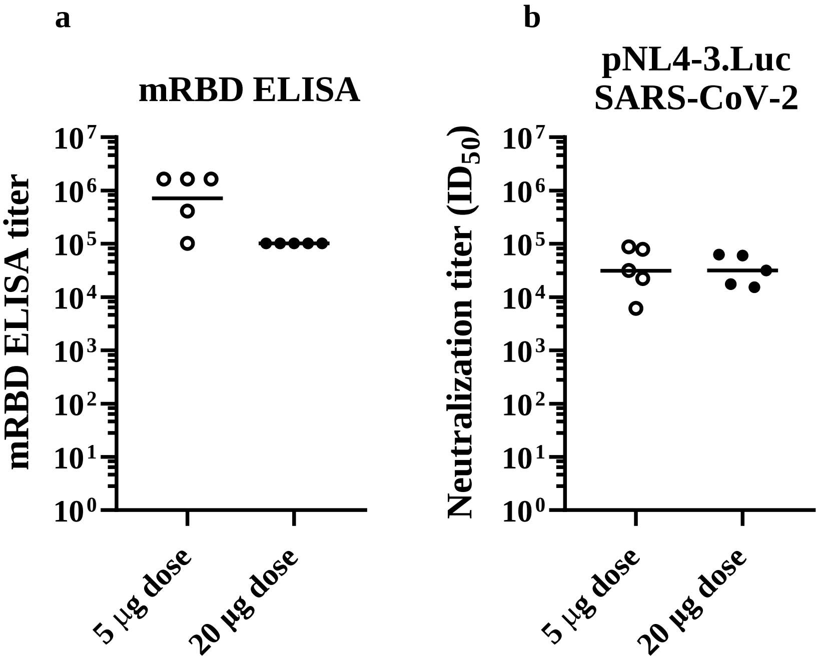


**Supporting Figure S9. ELISA titers with 5 µg of hCMP-mRBD dose. a.** Mice (n = 5/group) were immunized at week 0 and 3 with 5 µg of hCMP-mRBD adjuvanted with AddaVax™. Day 14 sera post boost was assayed for **A.** ELISA binding titers to mRBD. **b.** Pseudoviral neutralization titer utilizing pNL4-3.Luc. SARS-CoV-2 D614G Δ19. The black horizontal lines in each scatter plot represent Geometric mean titer (GMT).

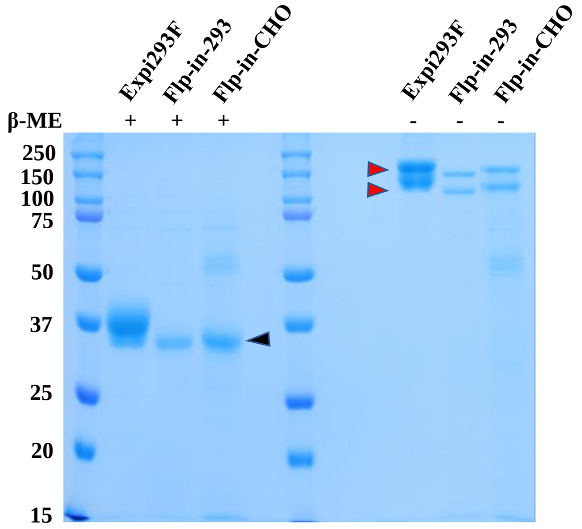


**Supporting Figure S10.** SDS-PAGE of purified hCMP-mRBD in reducing and non-reducing conditions. Protein was purified from transiently transfected *Expi293F* and stable cell lines Flp-in-293 and Flp-in-CHO**.** The black and red arrows represent the reduced and non-reduced protein bands respectively. The two red arrows likely indicate variably glycosylated forms.
